# Sequential co-translational assembly of the yeast SAGA DUB module precedes post-translational core docking

**DOI:** 10.64898/2026.11.15.766793

**Authors:** Andrea Barahona-Mejías, Jorge Abraham Haché-Santana, Lucía Muñoz-Alcantarilla, Alberto Elías-Villalobos

**Author notes:** corresponding author: Alberto Elías-Villalobos.

## Abstract

The 1.8-megadalton SAGA coactivator is organized into discrete functional modules, yet how individual catalytic modules are assembled and integrated into the holo-complex remains poorly understood. Here, we dissect the assembly pathway of the four-subunit histone H2B deubiquitinating (DUB) module in *Saccharomyces cerevisiae*. Pairwise RIP-qPCR uncovers a unidirectional network of co-translational interactions sequentially linking Sus1, Sgf11, Ubp8, and Sgf73. Combining systematic deletion genetics with co-immunoprecipitation and quantitative NanoBiT assays in intact living cells, we demonstrate that these contacts form an obligate linear cascade rather than independent binary events: each upstream subcomplex licenses the next nascent-chain engagement, sustains downstream Sgf11 and Ubp8 stability, and precludes post-translational bypass. Conversely, DUB subunits exhibit no co-translational interactions with the SAGA core interface, indicating that completed modules dock post-translationally. Thus, unlike single-scaffold assembly models, SAGA separates multi-step co-translational module templating from post-translational holo-complex docking, safeguarding compositional fidelity.

## Introduction

Assembly of multisubunit protein complexes is a tightly coordinated process that ensures correct folding, stoichiometry, and functional integrity, achieved through general and dedicated chaperone systems (1–4) and, in some cases, spatial compartmentalization (5). In addition, co-translational assembly has emerged as a key and evolutionarily conserved principle for protein complex biogenesis in eukaryotes, whereby subunits engage one another during translation through either simultaneous interactions between nascent chains or sequential binding of a fully synthesized subunit to a nascent partner (6–17). Beyond promoting complex formation, these co-translational interactions can also stabilize the nascent subunit itself, protecting it from misfolding and degradation (17–19). By constraining the timing, order, and directionality of subunit engagement, co-translational assembly therefore raises a fundamental question: does it primarily enhance the efficiency and fidelity of complex biogenesis, or does it constitute an obligate mechanism that dictates the final architecture of the complex? Although current evidence indicates that co-translational assembly can serve both functions (for recent review see (20)), the successful *in vitro* reconstitution of co-translationally assembling complexes from individually expressed subunits, leaves open the question of whether co-translational assembly can be bypassed through post-translational binding of subunits or if it is an obligate mechanism for protein complex formation in living cells. For instance, in the yeast nuclear pore complex, the Nsp1-Nup57 pair assembles co-translationally *in vivo* (12,14), yet associate spontaneously *in vitro* when independently expressed in *Escherichia coli* and subsequently mixed (21). Additionally, evolutionarily conserved complexes can differ, even minimally, in subunit composition or interacting partners, which may be reflected in the co-or post-translational assembly of their subunits or the order of subunit incorporation.

The SAGA complex offers a well-suited model to address these questions. This evolutionarily conserved transcriptional co-activator plays a prominent role in the regulation of inducible genes (22), with relevance across physiological and pathological contexts (23–29). In yeast, SAGA comprises 19 subunits (20 in humans) organized into discrete functional modules (30): a structural core; a TBP-binding module; a transcription factor-interacting module built around Tra1 (TRRAP in humans); and two enzymatic modules, with histone acetyltransferase (HAT) and histone deubiquitinase (DUB) activities. The structure of the SAGA core has been resolved in several organisms, including *Saccharomyces cerevisiae*, *Chaetomium thermophilum*, and *Komagataella phaffii* (31–33), and is conserved among these fungi, but differs in humans (34) – despite yeast and human SAGA sharing 18 of their 19-20 subunits. This raises the possibility that achieving divergent architectures from largely equivalent sets of subunits requires distinct assembly pathways, making SAGA a particularly informative system in which to study the relationship between assembly and structural diversification.

Yet, how SAGA assembles – particularly at the co-translational level – remains largely unresolved. Only a limited number of pairwise co-translational interactions have been reported to date, involving subunits from the HAT, core and DUB modules (10,35,36). In yeast, the HAT subunit Ada2 binds nascent Spt20 (35). In human cells, three pairwise co-translational interactions have been reported for the core module – TAF9 with nascent TAF6L, TAF12 with nascent TADA1, and TAF5 with nascent SUPT20H (36) – and the DUB subunit ENY2 (yeast Sus1) interacts with nascent ATXN7L3 (Sgf11) (10). Whether these interactions occur sequentially or simultaneously, whether additional subunits engage co-translationally, and whether stable assembly intermediates exist remain unknown. Consequently, an integrated model of SAGA assembly, at the level of individual modules and of the full complex, is lacking.

This gap is particularly consequential for the DUB module. The HAT module and the core TAF subunits are always found within a stable assembly – SAGA itself or a closely related complex such as ADA/ATAC and TFIID (30) – and appear poorly tolerated outside one: in human cells, disruption of the SAGA core disengages the HAT module and leaves its catalytic subunit KAT2A uncomplexed, whereupon it is targeted for proteasomal degradation by the orphan quality control machinery (37). The DUB module behaves differently: in humans, ATXN7L3 and ENY2 associate not only with USP22 within SAGA but also with the USP22-related proteases USP27X and USP51, forming complexes that lack ATXN7, the subunit anchoring the module to SAGA (38). Similarly, plants lack an ATXN7/Sgf73 ortholog altogether, yet their remaining DUB subunits remain active outside the context of SAGA (39,40). Rather than being degraded when not incorporated into SAGA, the DUB module thus appears able to persist and remain functional in its absence. Whether these independent forms arise through a dedicated biogenesis pathway or through remodeling of an assembled complex remains an open question. In yeast, the proteasome regulatory particle has been shown to actively dissociate a catalytically competent Sgf73-Ubp8 complex from assembled SAGA (41), demonstrating that remodeling constitutes at least one route to DUB independence – though whether a comparable outcome can also be reached directly, without prior incorporation into SAGA, remains unknown. Determining how the DUB module assembles may therefore help resolve this question and clarify how assembly pathways shape the molecular fate of SAGA’s modules.

Here, we investigate the assembly of the yeast SAGA DUB module, comprising Sus1, Sgf11, Ubp8, and Sgf73. We show that its biogenesis is driven by sequential, unidirectional co-translational interactions: Sus1 engages nascent Sgf11, which subsequently engages nascent Ubp8, which in turn engages nascent Sgf73, establishing a linear and staggered assembly pathway. Moreover, early co-translational engagements sustain the cellular stability of downstream module partners. Using yeast genetics and systematic *in vivo* analyses of wild-type and mutant strains, we validate this pathway and further show that the DUB module assembles as a discrete intermediate that is incorporated into SAGA post-translationally. These findings indicate that, for the DUB module, co-translational assembly constitutes an obligate, strictly ordered pathway required for its biogenesis *in vivo*. Collectively, these discoveries establish a coherent, stepwise model for SAGA DUB assembly and underscore the central role of assembly pathways in shaping the organization and function of multisubunit transcriptional regulators.

## Results

### Sus1 interacts co-translationally with nascent DUB subunits in a unidirectional manner

To evaluate whether co-translational assembly of the SAGA DUB module is evolutionarily conserved and extends beyond individual subunit pairs, we systematically examined the interaction network of *S. cerevisiae* Sus1. While human ENY2 (the Sus1 ortholog) is known to interact co-translationally with ATXN7L3, sequence divergence – such as the large C-terminal extension in ATXN7L3 relative to yeast Sgf11 ((42) and Suppl. Fig S1A,B) – and metazoan-specific interactors (38) raise the question of whether this biogenesis mechanism is conserved and how it scales across the entire module.

We first tested whether fully synthesized Sus1 associates with *SGF11* mRNA. RNA immunoprecipitation followed by quantitative PCR (RIP-qPCR) using Sus1-HA3-TAP2 as bait revealed significant enrichment of *SGF11* mRNA relative to negative controls, including the DUB-unrelated *VTC3* mRNA and an untagged strain (Fig 1A). *SUS1* mRNA served as an internal positive control (Suppl. Fig S1C). Treatment with puromycin – which causes premature nascent chain release from ribosomes – abolished *SGF11* mRNA enrichment without altering total Sus1 protein levels in input lysates or its IP recovery efficiency (Fig 1A; Suppl. Fig S1D), confirming that the Sus1-*SGF11* association is translation-dependent. Furthermore, epitope tagging of Sus1 did not alter cellular growth under stress conditions known to impair *sus1*Δ mutants (Suppl. Fig S1E), supporting the functional integrity of the tagged protein.

**Figure 1.**
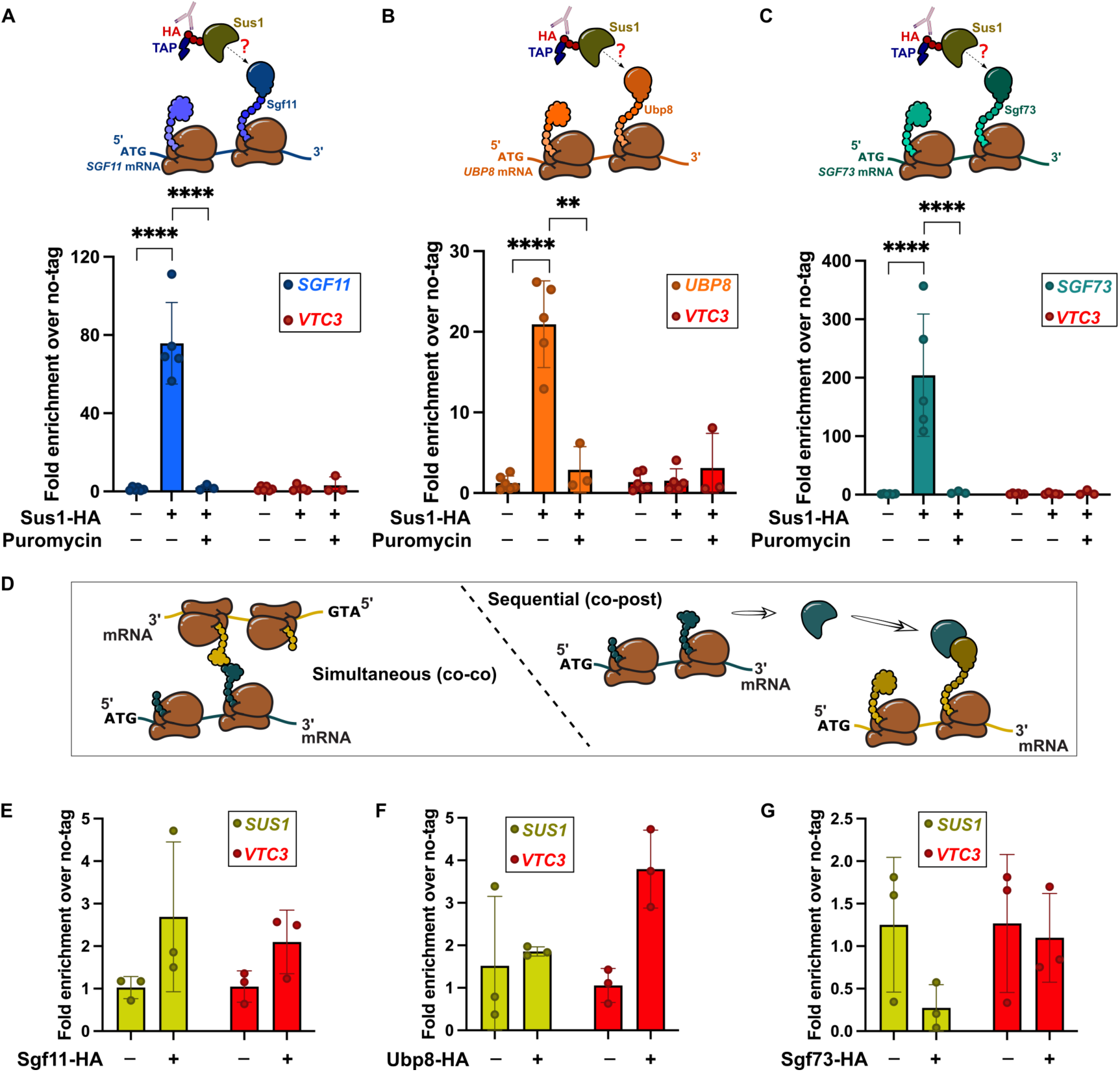
Sus1 unidirectionally engages nascent Sgf11, Ubp8, and Sgf73 during translation of the yeast SAGA DUB module. (A–C) Anti-HA RIP-qPCR analysis of *SGF11* (A), *UBP8* (B), and *SGF73* (C) mRNA enrichment alongside the unrelated negative control *VTC3* in untagged (−), Sus1-HA3-TAP2-tagged (+), and puromycin-treated Sus1-HA3-TAP2 (+ Puromycin) strains. Top diagrams illustrate the experimental setup testing whether immunoprecipitated Sus1-HA (pulled down via its HA3-TAP2 tag) associates with nascent Sgf11 (A), Ubp8 (B), or Sgf73 (C) polypeptides emerging from translating ribosomes. **(D)** Schematic model contrasting two modes of co-translational complex assembly: *simultaneous (co-co)* assembly (left), in which two nascent subunits interact concurrently while both remain ribosome-bound, versus *sequential (co-post)* assembly (right), in which a fully synthesized, ribosome-released subunit engages the nascent polypeptide chain of its partner on the ribosome. **(E–G)** Reciprocal anti-HA RIP-qPCR analysis of *SUS1* mRNA enrichment alongside *VTC3* in untagged (−) and Sgf11-HA3-TAP2 (E), Ubp8-HA3-TAP2 (F), or Sgf73-HA3-TAP2 (G) tagged (+) strains. In all graphs (A–C, E–G), individual data points represent independent biological replicates (*n* = 3–6), with mean ± SD overlaid, expressed as fold enrichment (2^-ΔΔCt^) relative to the mean Δ*Ct* of the untagged control. Individual replicates of this control are plotted to illustrate baseline variability. Statistical comparisons shown for the target DUB transcripts in (A–C) (untagged vs. Sus1-HA, and Sus1-HA vs. Sus1-HA + puromycin) were determined by two-way ANOVA on log_2_-transformed relative enrichment values followed by Tukey’s multiple comparisons test, using a significance threshold of α = 0.01 (*p* < 0.01, \*\*\*\**p* < 0.0001; panel B, Sus1-HA vs. Sus1-HA + puromycin, *p* = 0.0036; all other significant comparisons, *p* < 0.0001). Statistical comparisons were omitted in (E–G), where *SUS1* mRNA enrichment remained at background levels comparable to *VTC3*; *VTC3* is plotted alongside in all panels as an unrelated negative control to illustrate baseline background.

Contrary to nascent ATXN7L3 binding, ENY2 immunoprecipitates fail to enrich the transcript encoding the catalytic subunit USP22 at polysomes (10). Given that Ubp8 and Sgf73 represent the yeast orthologs of USP22 and ATXN7, respectively, we sought to determine whether Sus1 interactions in *S. cerevisiae* are similarly restricted or extend across the entire module. Under the same experimental conditions, Sus1 pull-downs enriched not only *SGF11* but also *UBP8* and *SGF73* mRNAs in a puromycin-sensitive manner (Fig 1B,C). This demonstrates that, in yeast, Sus1 engages nascent chains of all three DUB partners, establishing a comprehensive co-translational network across the whole module.

Finally, to establish the directionality of these interactions as simultaneous (co-co) or sequential (co-post) (Fig 1D), we performed reciprocal RIP-qPCR assays using tagged Sgf11, Ubp8, or Sgf73 as baits and probing for *SUS1* mRNA. None of these subunits enriched *SUS1* transcript above background levels (Fig 1E-G; Suppl. Fig S2A-D). Together, these findings demonstrate that Sus1 engages nascent Sgf11, Ubp8, and Sgf73 in a unidirectional manner, establishing Sus1 as a primary driver of co-translational DUB module assembly.

### Sequential co-translational engagement of Sgf11 and Ubp8 establishes a directional assembly hierarchy

Following the demonstration that Sus1 engages co-translationally with all other nascent DUB subunits, we next investigated whether subsequent steps in DUB module biogenesis are similarly driven by co-translational events. RIP-qPCR assays using Sgf11 as bait demonstrated a significant, puromycin-sensitive enrichment of both *UBP8* and *SGF73* transcripts (Fig 2A,B; Suppl. Fig S2A,D). Conversely, reciprocal pull-downs using tagged Ubp8 or Sgf73 failed to enrich *SGF11* mRNA above background levels (Fig 2C,D; Suppl. Fig S2B-D).

**Figure 2.**
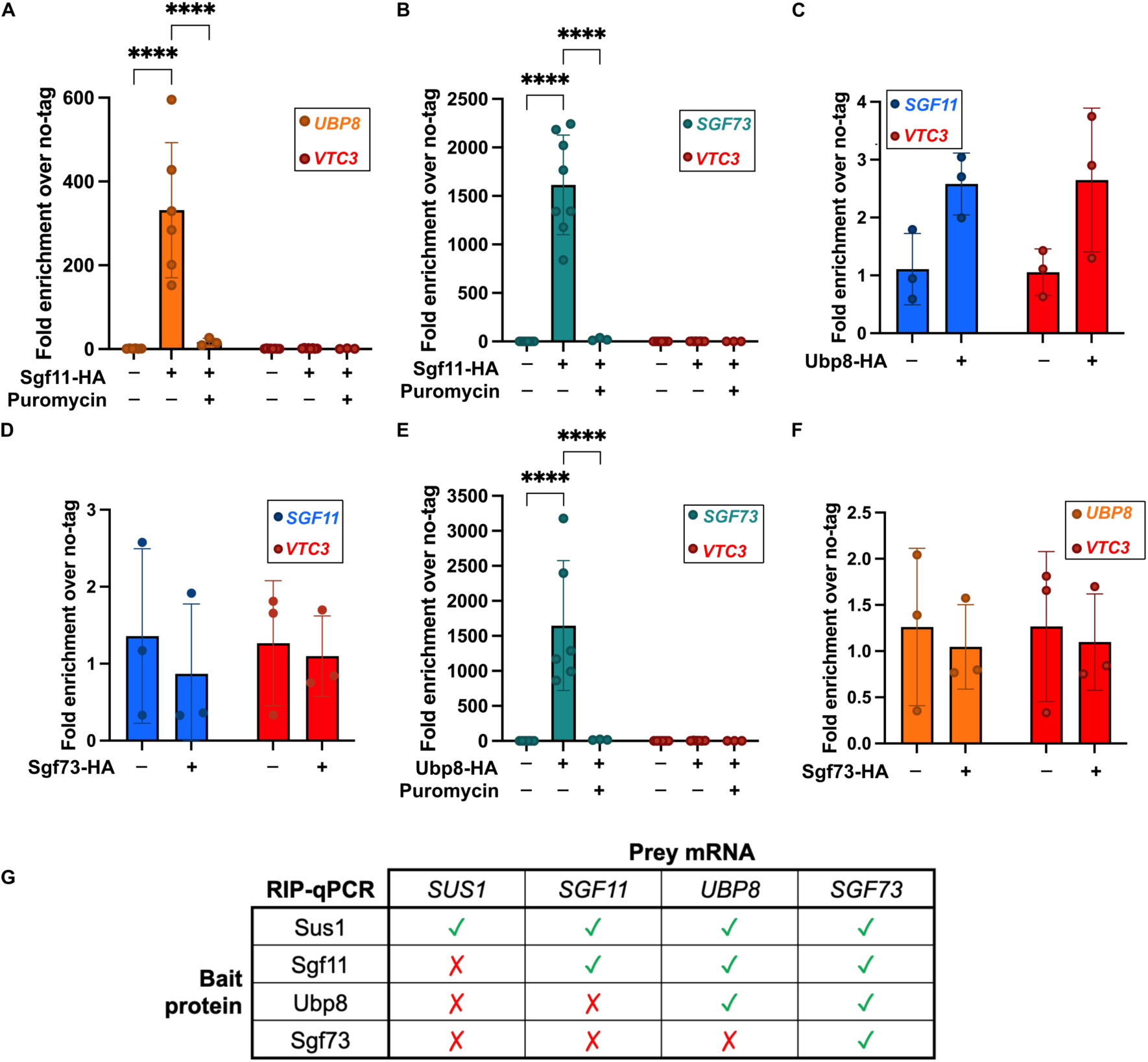
Co-translational assembly of the SAGA DUB module proceeds through a unidirectional cascade of pairwise subunit interactions. (A–F) Anti-HA RIP-qPCR analysis of target DUB subunit mRNA enrichment alongside the unrelated negative control *VTC3* in untagged (−), HA3-TAP2-tagged (+), and, where indicated, puromycin-treated (+ Puromycin) strains: Sgf11-HA3-TAP2 with *UBP8* (A) and *SGF73* (B) mRNAs; Ubp8-HA3-TAP2 with *SGF11* (C) and *SGF73* (E) mRNAs; and Sgf73-HA3-TAP2 with *SGF11* (D) and *UBP8* (F) mRNAs. Individual data points represent independent biological replicates (*n* = 3–9), with mean ± SD overlaid, expressed as fold enrichment (2^-ΔΔCt^) relative to the mean Δ*Ct* of the untagged control. Individual replicates of this control are plotted to illustrate baseline variability. Statistical comparisons shown for the target DUB transcripts in (A, B, E) (untagged vs. tagged, and tagged vs. tagged + puromycin) were determined by two-way ANOVA on log_2_-transformed relative enrichment values followed by Tukey’s multiple comparisons test, using a significance threshold of α = 0.01 (\*\*\*\**p* < 0.0001). Statistical comparisons were omitted in (C, D, F), where target mRNA enrichment remained at background levels comparable to *VTC3*; *VTC3* is plotted alongside in all panels as an unrelated negative control to illustrate baseline background. **(G)** Summary matrix of all pairwise RIP-qPCR interactions between immunoprecipitated DUB bait proteins (rows) and prey mRNAs (columns), integrating the heteromeric interactions from Figures 1 and 2 together with self-mRNA controls (Supplementary Figs. S1C and S2). Green checkmarks (✓) indicate specific puromycin-sensitive co-translational enrichment, whereas red crosses (✗) indicate absence of enrichment above background.

To complete the pairwise mapping of co-translational contacts within the DUB module, we evaluated the interaction between Ubp8 and Sgf73. Immunoprecipitation of Ubp8 enriched *SGF73* mRNA in a puromycin-sensitive manner, whereas reciprocal Sgf73 pull-downs showed no enrichment of the *UBP8* transcript (Fig 2E,F; Suppl. Fig S2B-D), establishing a unidirectional interaction from Ubp8 toward nascent Sgf73.

Together, these reciprocal RIP assays establish a comprehensive, fully mapped matrix of pairwise unidirectional contacts within the module: Sus1 engages nascent Sgf11, Ubp8, and Sgf73; Sgf11 engages nascent Ubp8 and Sgf73; and Ubp8 engages nascent Sgf73, with no reverse co-translational binding detected in any instance (Fig 2G). Rather than independent or stochastic binding events, the absolute directionality maintained across every interacting pair strongly implies that co-translational DUB module assembly follows a strictly governed, step-wise pathway.

### Upstream incorporated DUB subunits sustain stability of downstream engaged partners

Co-translational interactions often fulfill a dual role, facilitating multi-protein complex assembly while simultaneously guiding the folding and stabilization of nascent polypeptides (17). To determine whether individual DUB subunits are required to maintain the stability of their module partners under our experimental conditions, we systematically analyzed the steady-state protein abundance of each DUB component in single deletion strains by Western blotting. If inter-subunit stability dependencies govern module integrity, our unidirectional co-translational assembly model predicts that loss of a given subunit will selectively compromise the stability of downstream, but not upstream, partners.

In full agreement with this hypothesis, Western blot analyses revealed that Sus1 protein levels remained unaffected across *sgf11*Δ, *ubp8*Δ, and *sgf73*Δ strains (Fig 3A-C). RT-qPCR measurements confirmed that *SUS1* mRNA abundance was similarly unreduced in all three mutant backgrounds (Suppl. Fig S3A–C). This sustained steady-state level of Sus1 protein aligns with independent reports showing that Sus1 expression and stability are independent of *SGF11* (41), *UBP8*, or *SGF73* (42).

**Figure 3.**
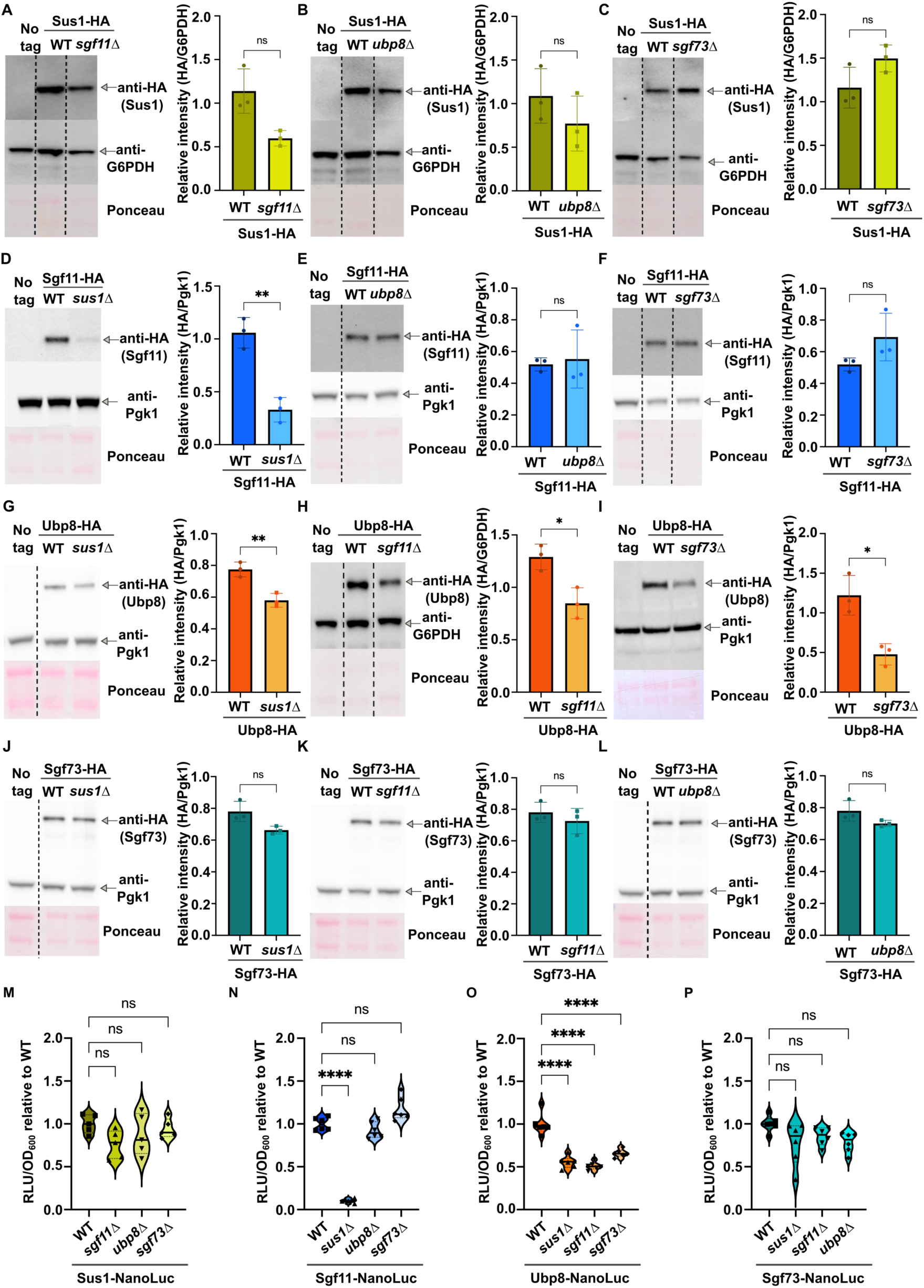
DUB module subunits exhibit hierarchical steady-state stability dependencies in yeast. (A–L) Western blot analysis of steady-state protein levels of HA3-TAP2-tagged DUB module subunits in wild-type (WT) and the indicated deletion mutant strains, alongside untagged controls (No tag): Sus1-HA in WT versus *sgf11*Δ (A; *p* = 0.0537), *ubp8*Δ (B; *p* = 0.2833), and *sgf73*Δ (C; *p* = 0.1184); Sgf11-HA in WT versus *sus1*Δ (D; *p* = 0.0029), *ubp8*Δ (E; *p* = 0.7843), and *sgf73*Δ (F; *p* = 0.1776); Ubp8-HA in WT versus *sus1*Δ (G; *p* = 0.0062), *sgf11*Δ (H; *p* = 0.0173), and *sgf73*Δ (I; *p* = 0.0191); and Sgf73-HA in WT versus *sus1*Δ (J; *p* = 0.0706), *sgf11*Δ (K; *p* = 0.4083), and *ubp8*Δ (L; *p* = 0.1559). In each panel, representative anti-HA immunoblots are shown on the left alongside the corresponding loading control—G6PDH for Sus1-HA (A–C) and Ubp8-HA in *sgf11*Δ (H), and Pgk1 for all remaining panels (D–G, I–L)—and Ponceau S staining to verify total protein transfer. Vertical dashed lines indicate where intervening lanes from the same membrane were spliced out. Densitometric quantifications are shown on the right as the ratio of HA signal to the corresponding loading control; individual data points represent independent biological replicates (*n* = 3), with mean ± SD overlaid. Statistical significance was determined by a two-tailed unpaired Welch’s *t*-test with α = 0.05 (\**p* < 0.05; \*\**p* < 0.01; ns, not significant, *p* ≥ 0.05). **(M–P)** Luminescence-based quantification (RLU/OD_600_ normalized to the mean of the corresponding WT control) of steady-state levels of C-terminally NanoLuc-tagged DUB subunits in WT and the indicated deletion mutants: Sus1-NanoLuc in WT, *sgf11*Δ (*p* = 0.0886), *ubp8*Δ (*p* = 0.4929), and *sgf73*Δ (*p* = 0.9062) (M; *n* = 5); Sgf11-NanoLuc in WT, *sus1*Δ (*p* < 0.0001), *ubp8*Δ (*p* = 0.3645), and *sgf73*Δ (*p* = 0.124) (N; *n* = 6); Ubp8-NanoLuc in WT, *sus1*Δ (*p* < 0.0001), *sgf11*Δ (*p* < 0.0001), and *sgf73*Δ (*p* < 0.0001) (O; *n* = 6); and Sgf73-NanoLuc in WT, *sus1*Δ (*p* = 0.0524), *sgf11*Δ (*p* = 0.2636), and *ubp8*Δ (*p* = 0.0654) (P; *n* = 6). Violin plots show the distribution of individual biological replicates, with horizontal lines indicating the median and quartiles. Statistical significance relative to WT was determined by one-way ANOVA followed by Dunnett’s post-hoc test using a significance threshold of α = 0.01 (\*\*\*\**p* < 0.0001; ns, not significant, *p* ≥ 0.01).

In stark contrast to Sus1, Sgf11 protein abundance was severely compromised upon *SUS1* deletion, displaying a nearly 70% reduction relative to wild-type levels (Fig 3D). This marked decline mirrors the structural interdependence reported for their metazoan orthologs, where ENY2 is strictly required for ATXN7L3 stabilization (43). Conversely, Sgf11 protein levels remained at wild-type amounts in *ubp8*Δ, and *sgf73*Δ strains (Fig 3E,F). As observed for *SUS1*, *SGF11* transcript levels were not negatively altered in any mutant background (Suppl. Fig S3D-F), confirming that the marked reduction of Sgf11 protein in *sus1*Δ cells is not driven by decreased transcript abundance.

Regarding Ubp8, Western blot analyses revealed a moderate reduction in total protein abundance in *sus1*Δ, *sgf11*Δ, and *sgf73*Δ strains, whereas *UBP8* mRNA levels did not decrease in any mutant (Fig 3G-I; Suppl. Fig S3G-I). The decline of Ubp8 in *sus1*Δ and *sgf11*Δ backgrounds aligns with the post-translational dependency reported in human cells, where ATXN7L3 is required for USP22 accumulation (40), supporting a role for upstream co-translational assembly partners in safeguarding downstream subunit stability. Furthermore, the moderate decrease of Ubp8 observed in *sgf73*Δ cells indicates that while Sgf73 is not strictly required for initial DUB heterotrimer formation, loss of Sgf73 impacts overall Ubp8 accumulation, consistent with its structural requirement for Ubp8 catalytic domain integrity (44).

Finally, steady-state Sgf73 protein levels were completely unaffected by the deletion of any of the other three DUB subunits (Fig 3J-L; Suppl. Fig S3J-L), indicating that co-translational engagement by upstream subunits plays little to no role in Sgf73 stability. This resilience is consistent with biochemical evidence showing that orphan Sgf73 can independently incorporate into the SAGA core scaffold in the absence of the other DUB components (41,44), confirming that uncomplexed Sgf73 remains structurally competent for SAGA binding.

Although these steady-state analyses support a model where upstream assembly steps (Sus1 → Sgf11 → Ubp8) protect downstream partners from degradation, the persistence of a substantial Ubp8 pool across deletion mutants, as well as the wild-type stability of Sgf73, could theoretically reflect altered protein turnover kinetics. To test whether accelerated degradation was masking a more pronounced instability, we performed cycloheximide chase assays for Ubp8 and Sgf73 in *sus1*Δ cells. These experiments revealed comparable degradation kinetics for both Ubp8 and Sgf73 in *sus1*Δ relative to wild-type strains (Suppl. Fig S4), ruling out increased turnover as the primary determinant of their persistence.

To complement Western blotting with an orthogonal, highly quantitative *in situ* readout, we developed a luminescence-based reporter assay using NanoLuc luciferase fused to individual DUB subunits in permeabilized cells. As shown in Fig 3M-P, the NanoLuc approach faithfully recapitulated our Western blot findings with precise quantitative resolution. Deletion of *SUS1* caused an 88% drop in Sgf11-NanoLuc signal (Fig 3N). Similarly, Ubp8-NanoLuc luminescence exhibited moderate reductions in *sus1*Δ (43%), *sgf11*Δ (47%), and *sgf73*Δ (30%) mutants (Fig 3O). All other subunit-NanoLuc combinations across the respective deletion backgrounds maintained luminescence levels comparable to wild-type controls (Fig 3M–P).

Altogether, these systematic protein-level analyses confirm that subunits positioned upstream in the co-translational assembly pathway are essential for the full stabilization of downstream partners. The sole exception is the terminal subunit, Sgf73, whose stability is intrinsically self-sustained, restricting its role to licensing the incorporation of the fully assembled DUB module into the SAGA core complex (45).

### Genetic dissection confirms a strictly linear hierarchy of co-translational DUB module assembly

Having established that upstream DUB components sustain the post-translational stability of downstream partners, we next investigated whether this directional dependency reflects an obligate co-translational assembly cascade during translation (Sus1 → Sgf11 → Ubp8 → Sgf73; Fig 4A). To test whether upstream subunit incorporation is required to license the co-translational engagement of downstream nascent chains – rather than merely influencing mature protein abundance – we systematically performed RIP-qPCR assays across single DUB deletion strains. Under an obligate linear assembly hierarchy, the loss of an upstream anchor is predicted to selectively abolish co-translational engagement of downstream nascent polypeptides, while leaving upstream nascent-chain engagements unperturbed.

**Figure 4.**
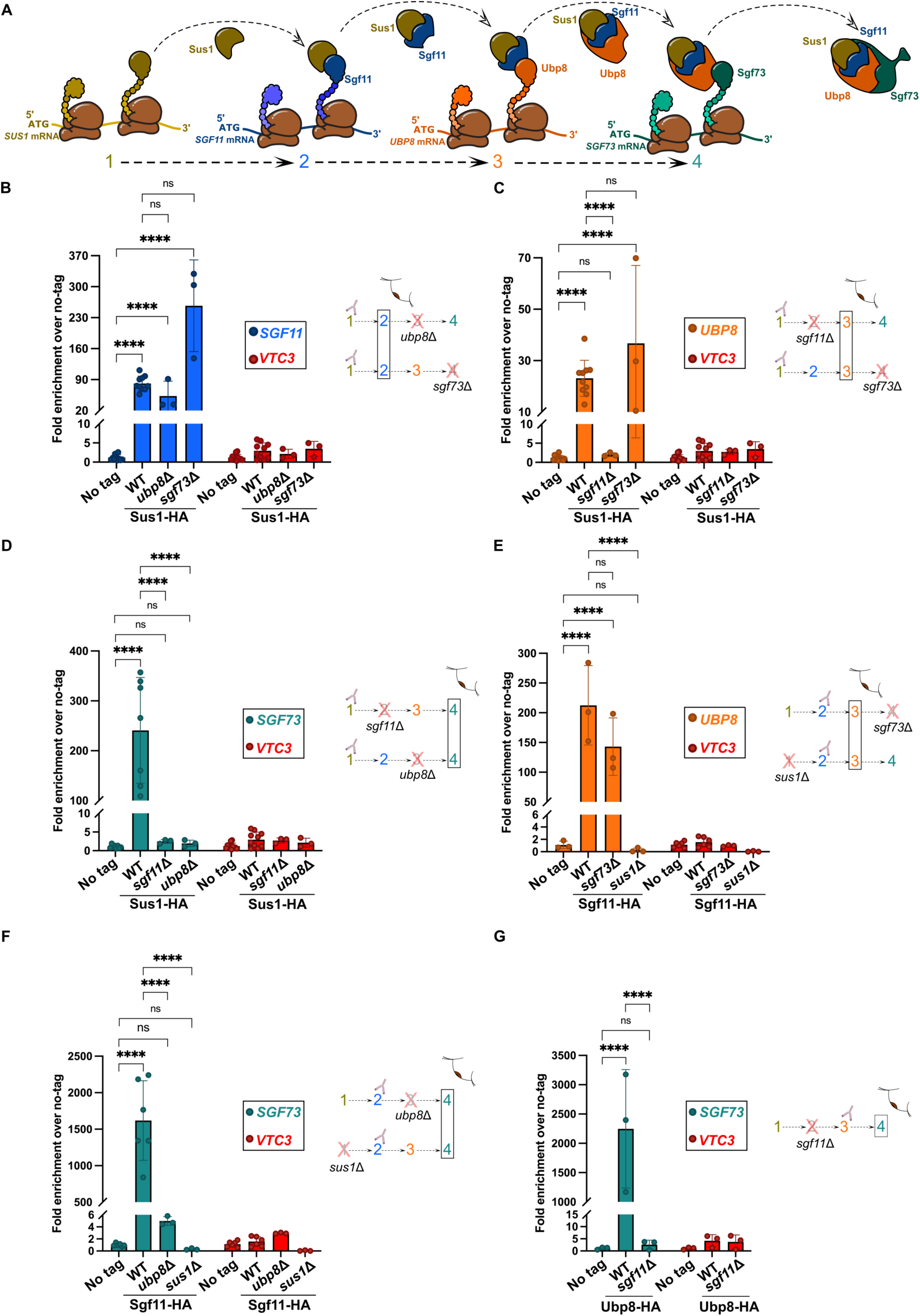
Co-translational assembly of the yeast SAGA DUB module follows a strict hierarchical pathway. **(A)** Working model illustrating the ordered, directional co-translational assembly pathway of the DUB module (steps 1–4): newly synthesized Sus1 (1) co-translationally engages nascent Sgf11 (2); the resulting Sus1–Sgf11 subcomplex co-translationally recruits nascent Ubp8 (3); and the pre-assembled Sus1–Sgf11–Ubp8 trimer subsequently engages nascent Sgf73 (4) to complete DUB module assembly. **(B–G)** Anti-HA RIP-qPCR analysis of target DUB subunit mRNA enrichment alongside the unrelated negative control *VTC3* in untagged (No tag), wild-type (WT), and the indicated deletion mutant backgrounds: Sus1-HA3-TAP2 immunoprecipitations quantifying *SGF11* (B), *UBP8* (C), and *SGF73* (D) mRNAs; Sgf11-HA3-TAP2 immunoprecipitations quantifying *UBP8* (E) and *SGF73* (F) mRNAs; and Ubp8-HA3-TAP2 immunoprecipitations quantifying *SGF73* mRNA (G). Diagrams to the right of each graph reference steps 1-4 in (A), indicating the immunoprecipitated HA-tagged bait (antibody symbol), deleted subunits (red crosses), and the quantified target mRNA (boxed step). Individual data points represent independent biological replicates (*n* = 3–11), with mean ± SD overlaid, expressed as fold enrichment (2^-ΔΔCt^) relative to the mean Δ*Ct* of the untagged control. Individual replicates of this control are plotted to illustrate baseline variability. Statistical comparisons shown for the target DUB transcripts were determined across experimental conditions by two-way ANOVA on log_2_-transformed relative enrichment values followed by Tukey’s multiple comparisons test (simple effects within each transcript), using a significance threshold of α = 0.01 (\*\*\*\**p* < 0.0001; ns, not significant, *p* ≥ 0.01); *VTC3* is plotted alongside as an unrelated negative control to illustrate baseline background.

Evaluating Sus1 as the primary upstream anchor fully validated these predictions. Sus1 co-translational engagement with *SGF11* mRNA remained intact in both *ubp8*Δ and *sgf73*Δ strains (Fig 4B). In contrast, Sus1 binding to the *UBP8* transcript was abolished in *sgf11*Δ cells but preserved in *sgf73*Δ mutants (Fig 4C). Furthermore, Sus1 association with *SGF73* mRNA was lost upon deletion of either *SGF11* or *UBP8* (Fig 4D). As an internal positive control for immunoprecipitation efficiency and nascent chain recovery, Sus1 robustly enriched its own mRNA (*SUS1*) across all deletion backgrounds (Suppl. Fig S5A), with equivalent protein recovery across strains confirmed by Western blot (Suppl. Fig S5D).

Further confirming this linear dependency downstream, Sgf11 RIP-qPCR assays showed that *SGF73* deletion had no effect on the ability of Sgf11 to engage the *UBP8* transcript (Fig 4E). Conversely, *UBP8* deletion abolished co-translational enrichment of *SGF73* mRNA by Sgf11 (Fig 4F). Beyond these pairwise requirements, we tested whether an upstream anchor is required to license the binding capacity of downstream subunits – that is, whether Sgf11 must be previously engaged by Sus1 to recruit subsequent targets. Strikingly, *SUS1* deletion eliminated Sgf11 association with both *UBP8* and *SGF73* mRNAs (Fig 4E,F). Importantly, Sgf11 retained its capacity to enrich its own transcript (*SGF11*) in *sus1*Δ cells (Suppl. Fig S5B,D), serving as an internal positive control for the recovery of translating Sgf11 nascent chains despite reduced overall protein abundance and IP recovery (Fig 3D,N; Suppl. Fig S5D). This confirms that the assay successfully captured translating Sgf11, indicating that its failure to recruit downstream nascent chains stems from its rapid post-translational clearance or its structural inability to function as a binding platform in the absence of Sus1.

This requirement for an upstream structural anchor during sequential assembly is most unequivocally demonstrated by Ubp8 pull-downs in the *sgf11*Δ background. Although Ubp8 protein levels are only modestly affected by the loss of *SGF11* (Fig 3H,O), its co-translational interaction with *SGF73* mRNA completely disappeared in *sgf11*Δ cells (Fig 4G). Ubp8 retained robust enrichment of its own transcript (*UBP8*) in *sgf11*Δ cells (Suppl. Fig S5C,D), ruling out technical or expression artifacts. This demonstrates that without prior engagement by Sgf11, Ubp8 is intrinsically incapable of recruiting nascent Sgf73, proving that downstream subunits cannot act as isolated anchors but must assemble as a pre-formed upstream platform to license subsequent assembly steps.

Together, these results provide definitive genetic proof that DUB module biogenesis follows an obligate, unidirectional, and strictly hierarchical co-translational assembly pathway.

### In vivo physical interaction networks reflect DUB module assembly hierarchy

While genetic dissection by RIP-qPCR establishes an obligate co-translational assembly pathway, it does not inform *per se* on the post-translational fate, physical persistence, or structural integrity of the resulting mature subcomplexes. Crucially, it remains unclear whether subcomplexes formed co-translationally maintain their physical assembly post-translationally, or whether subunits can associate spontaneously post-translationally if co-translational biogenesis is bypassed. To address this, we performed systematic co-immunoprecipitation (Co-IP) assays evaluating every pairwise subunit interaction across wild-type (WT) and single DUB deletion strains in soluble cell extracts.

In complete agreement with our co-translational assembly model (Sus1 → Sgf11 → Ubp8 → Sgf73), physical association between mature proteins faithfully recapitulated the co-translationally established hierarchy. Analysis of Sus1-centered complexes showed that Sus1 co-immunoprecipitated Sgf11 in wild-type, *ubp8*Δ, and *sgf73*Δ strains, demonstrating that the Sus1-Sgf11 physical association persists post-translationally, mirroring their co-translational engagement independently of downstream subunits (Fig. 5A). In contrast, the mature Sus1-Ubp8 physical interaction was lost in *sgf11*Δ cells but remained preserved in *sgf73*Δ strains (Fig 5B), while Sus1 failed to associate post-translationally with Sgf73 upon deletion of either *SGF11* or *UBP8* (Fig 5C). Examining Sgf11-containing subcomplexes, Sgf11 maintained its physical association with Ubp8 in *sgf73*Δ cells while the remaining Sgf11 protein in *sus1*Δ mutants (Fig 3D,N) lost its binding to Ubp8 (Fig 5D). On its side, the Sgf11-Sgf73 physical interaction fell below detection limits in both *sus1*Δ and *ubp8*Δ backgrounds (Fig 5E). Finally, Ubp8 failed to co-immunoprecipitate Sgf73 in *sus1*Δ and *sgf11*Δ strains (Fig 5F).

**Figure 5.**
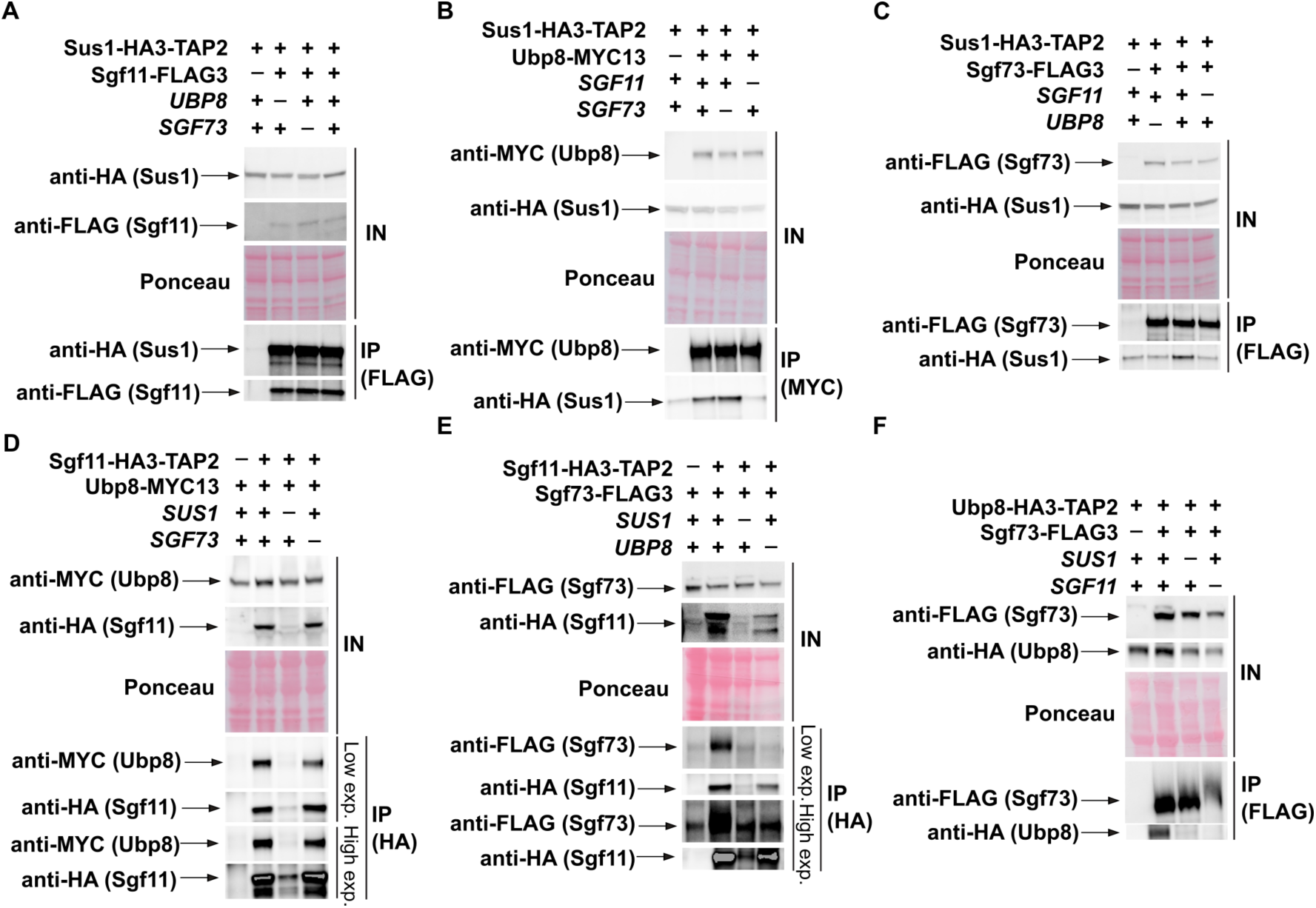
Stable pairwise physical interactions within the DUB module require upstream assembly partners. (A–F) Co-immunoprecipitation (co-IP) assays examining pairwise physical interactions between tagged DUB module subunits in wild-type (+) and the indicated single-subunit deletion (−) backgrounds, alongside untagged control strains (−): Sus1-HA3-TAP2 with Sgf11-FLAG3 (A), Ubp8-MYC13 (B), or Sgf73-FLAG3 (C); Sgf11-HA3-TAP2 with Ubp8-MYC13 (D) or Sgf73-FLAG3 (E); and Ubp8-HA3-TAP2 with Sgf73-FLAG3 (F). Input (IN) and immunoprecipitated (IP, pulled down via the FLAG, MYC, or HA epitope as indicated in parentheses) fractions were analyzed by immunoblotting with anti-HA, anti-FLAG, and anti-MYC antibodies. Input membranes were stained with Ponceau S to verify protein loading and transfer. In panels (D) and (E), both low (Low exp.) and high (High exp.) exposures of the IP blots are shown to allow detection of residual co-immunoprecipitation when Sgf11 steady-state levels are reduced in the *sus1*Δ background.

The requirement of Sgf11 for the Sus1-Ubp8 interaction aligns with structural and biochemical models positioning Sgf11 as an essential physical scaffold for DUB module integrity (43,44), providing a mechanistic basis for classical observations showing that Ubp8 fails to integrate into SAGA in *sgf11*Δ mutants (45,46). Crucially, the finding that the core trimeric interaction between Sus1, Sgf11, and Ubp8 remains assembled in *sgf73*Δ cells ((46,47) and Fig 5A,B,D) confirms that subcomplexes generated co-translationally maintain their physical integrity post-translationally, even when detached from the SAGA core scaffold (48). Furthermore, the loss of downstream physical interactions in upstream deletion strains confirms that post-translational association cannot spontaneously rescue or bypass a failure in co-translational assembly.

Because Co-IPs, and RIPs, rely on cell lysis – which removes molecular interactions from their native cellular context – we next leveraged luminescence-based assays in living cells as an orthogonal approach to further validate these findings *in vivo*. To probe the spatial proximity and structural arrangement of the DUB module in living cells we used split-luciferase complementation technology (NanoBiT). Under physiological growth conditions (rich media, OD_600_∼0.8), live-cell NanoBiT signals recapitulated the assembly hierarchy predicted by our linear co-translational model. In full agreement with our physical interaction data, loss of upstream subunits caused downstream complementation pairs to drop to negative control baseline levels (Fig 6). Specifically, *SUS1* deletion reduced Sgf11-Ubp8, Sgf11-Sgf73, and Ubp8-Sgf73 NanoBiT signals to baseline control values (Fig 6D-F). Similarly, *SGF11* deletion abolished Sus1-Ubp8, Sus1-Sgf73 and Ubp8-Sgf73 complementation (Fig 6B,C,F). Conversely, the upstream Sus1-Sgf11 interaction remained at wild-type levels in both *ubp8*Δ and *sgf73*Δ strains (Fig 6A), reinforcing that early core interactions form independently of downstream module members. Crucially, NanoBiT analysis in *sgf73*Δ cells revealed a distinct structural nuance regarding the soluble DUB subcomplex. While Sgf11-Ubp8 complementation showed 2.25-fold increased luminescence in *sgf73*Δ cells as compared to the wild-type (Fig 6D), Sus1-Ubp8 complementation exhibited a pronounced quantitative reduction of ∼15-fold (Fig 6B). Importantly, despite this decrease relative to wild-type, the Sus1-Ubp8 signal in *sgf73*Δ cells remains ∼174-fold above its respective negative control baseline, in sharp contrast to *sgf11*Δ mutants where signal is completely lost (Figure 6B). Given that Sus1, Sgf11, and Ubp8 stay physically associated in *sgf73*Δ soluble extracts ((46,47) and Fig 5A,B,D), this specific attenuation indicates that *SGF73* deletion does not physically dismantle the Sus1-Ubp8 complex, but rather introduces conformational flexibility or increased spatial distance between the two subunits *in vivo*. Similarly, but with the opposite effect, loss of *SGF73* improves luminescence emission by the Sgf11-Ubp8 pair (Fig 6D).

**Figure 6.**
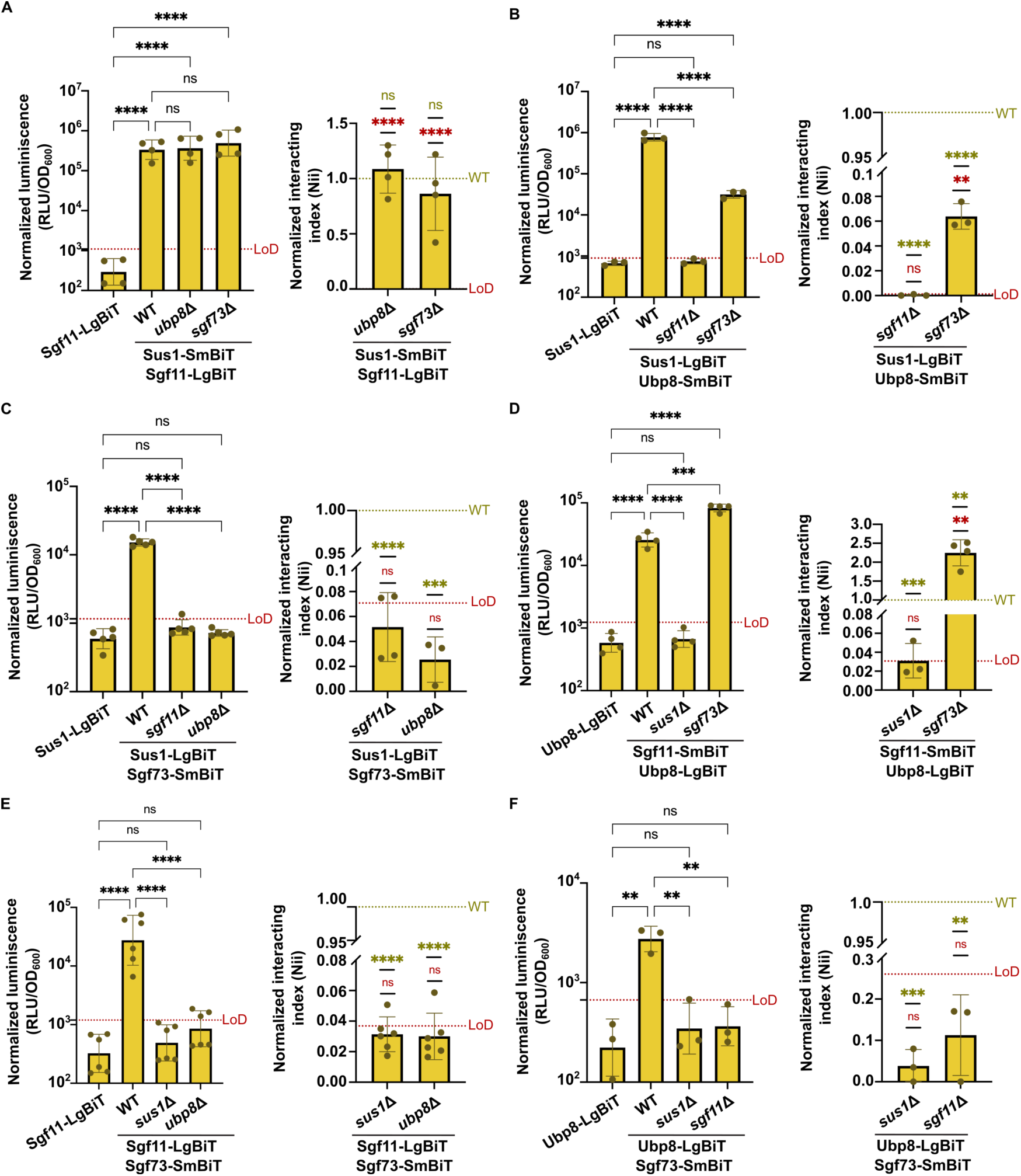
Live-cell interaction profiling confirms strict sequential assembly of the SAGA DUB module *in vivo*. (A–F) Quantitative NanoBiT protein–protein interaction assays measuring *in vivo* association between SmBiT- and LgBiT-tagged DUB module subunit pairs in wild-type (WT) and the indicated single-subunit deletion mutant backgrounds: Sus1-SmBiT/Sgf11-LgBiT (A), Sus1-LgBiT/Ubp8-SmBiT (B), Sus1-LgBiT/Sgf73-SmBiT (C), Sgf11-SmBiT/Ubp8-LgBiT (D), Sgf11-LgBiT/Sgf73-SmBiT (E), and Ubp8-LgBiT/Sgf73-SmBiT (F). In each panel, the left graph displays background-uncorrected luminescence normalized to culture optical density (RLU/OD_600_) alongside the corresponding single-subunit LgBiT-only negative control (lacking the SmBiT fusion); individual data points represent independent biological replicates, with mean ± SD overlaid, and the horizontal red dotted line indicates the limit of detection (LoD, defined as mean + 3 SD of the single-LgBiT negative control). The right graph shows the normalized interacting index (Nii) in the deletion mutants, calculated by subtracting the single-LgBiT background, normalizing to WT within each biological replicate (green dotted line at 1.00), and correcting for steady-state subunit abundance determined by NanoLuc assays (see Materials and Methods for details). Statistical comparisons in the left graphs were determined by one-way ANOVA on log_10_-transformed RLU/OD_600_ values followed by Šídák’s multiple comparisons test (α = 0.01). Statistical significance in the right graphs was evaluated on untransformed Nii values by two-tailed one-sample *t*-tests against a theoretical mean of 1.0 (WT, green asterisks) or against the LoD threshold (red asterisks) using α = 0.01 (*p* < 0.01, \*\*\**p* < 0.001, \*\*\*\**p* < 0.0001; ns, not significant, *p* ≥ 0.01 panel B right, *sgf73*Δ vs. LoD, *p* = 0.0089;; panel C right, *ubp8*Δ vs. WT, *p* = 0.0001; panel D left, WT vs. *sgf73*Δ, *p* = 0.0003; panel D right, *sus1*Δ vs. WT, *p* = 0.0001, *sgf73*Δ vs. WT, *p* = 0.0055, and *sgf73*Δ vs. LoD, *p* = 0.0010; panel F left, control vs. WT, *p* = 0.0017, WT vs. *sus1*Δ, *p* = 0.006, and WT vs. *sgf11*Δ, *p* = 0.007; panel F right, *sus1*Δ vs. WT, *p* = 0.0006, and *sgf11*Δ vs. WT, *p* = 0.004; all other significant comparisons, *p* < 0.0001).

Importantly, these findings reveal that the co-translational assembly pathway restricts the landscape of viable subcomplexes *in vivo*. Beyond verifying expected interactions, our analysis confirms the absence of theoretical intermediates that are excluded by the linear sequence. For instance, although structural contacts exist between Sgf11, Ubp8, and Sgf73 (43), a Sgf11-Ubp8-Sgf73 subcomplex fails to form in *sus1*Δ cells, and Ubp8 fails to associate with Sgf73 in *sgf11*Δ cells. This aligns with previous observations demonstrating that Ubp8 purifies as an isolated, unintegrated subunit in *sgf11*Δ mutants (46,47), and demonstrates that early co-translational engagement by upstream subunits is an absolute prerequisite to license the assembly of all downstream DUB components.

### The DUB module incorporates post-translationally into the SAGA core complex

Having established the linear co-translational pathway driving intra-module DUB assembly, we next investigated whether the fully formed DUB module incorporates into the broader SAGA core co-or post-translationally.

To test for potential co-translational cross-talk between the DUB module and the SAGA core, we targeted core subunits at the DUB binding interface. Cryo-EM maps demonstrate that the C-terminal region of Sgf73 serves as the primary anchor linking the DUB module to SAGA, extending into a core lobule formed by Spt20, the Ada1-Taf12 heterodimer, and Taf6-Taf9 ((31) and Fig 7A,B). Among these interface components, Spt20, Ada1, and Taf12 represent key structural candidates: Spt20 forms the main contact surface with Sgf73 (31) and its deletion abolishes DUB module integration into SAGA (49–51), while Ada1 and Taf12 constitute the core histone-fold backbone at this interaction site, with Taf6 and Taf9 showing rather peripheral contacts with Sgf73 (31).

**Figure 7.**
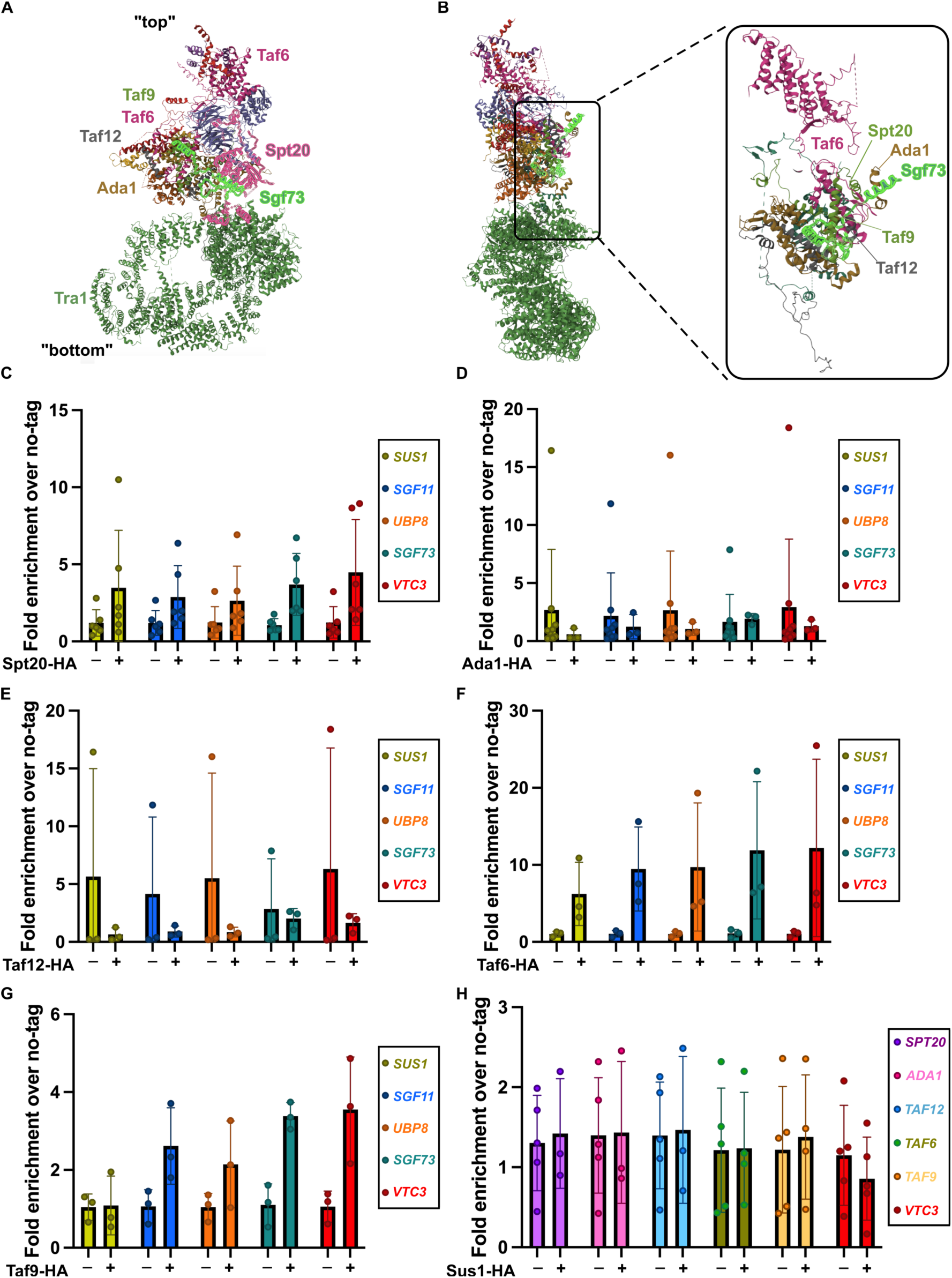
Co-translational assembly of the SAGA DUB module occurs independently of core SAGA complex subunits. (A, **B)** Front (A) and orthogonal (B) ribbon views of the *Saccharomyces cerevisiae* SAGA complex cryo-EM structure (PDB: 6T9I), showing the overall architecture from the Tra1 lobe (“bottom”) to the core module (“top”). The magnified inset in (B) highlights the structural interface where the DUB module anchor subunit Sgf73 (bright green) contacts neighboring SAGA core subunits (Spt20, Ada1, Taf6, Taf9, and Taf12). **(C–G)** Anti-HA RIP-qPCR analysis quantifying enrichment of the four DUB module mRNAs (*SUS1*, *SGF11*, *UBP8*, and *SGF73*) alongside the unrelated negative control *VTC3* in untagged (−) and HA-tagged (+) SAGA core subunit strains: Spt20-HA (C), Ada1-HA (D), Taf12-HA (E), Taf6-HA (F), and Taf9-HA (G). **(H)** Reciprocal anti-HA RIP-qPCR analysis in untagged (−) and Sus1-HA (+) strains quantifying enrichment of the core SAGA subunit mRNAs (*SPT20*, *ADA1*, *TAF12*, *TAF6*, and *TAF9*) alongside *VTC3*. In all graphs (C–H), individual data points represent independent biological replicates, with mean ± SD overlaid, expressed as fold enrichment (2^-ΔΔCt^) relative to the mean Δ*Ct* of the untagged control. Individual replicates of this control are plotted to illustrate baseline variability. Consistent with Figures 1E–G and 2C, D, and F, statistical comparisons were omitted because target mRNA enrichment remained at background levels comparable to *VTC3*, which is plotted alongside in all panels as an unrelated negative control to illustrate baseline background.

We therefore performed RIP-qPCR assays using Spt20, Ada1, Taf12, Taf6 and Taf9 to probe for the recruitment of nascent DUB module subunits. However, immunoprecipitation of any of these core subunits failed to detect any significant enrichment of *SUS1*, *SGF11*, *UBP8*, or *SGF73* mRNAs relative to the non-specific control transcript *VTC3* (Fig 7C-G; Suppl. Fig S6). To rule out potential detection limits in unidirectional assays, we conducted reciprocal RIP experiments by immunoprecipitating each individual DUB subunit (Sus1, Sgf11, Ubp8, or Sgf73) and quantifying the co-precipitation of *SPT20*, *ADA1*, *TAF12*, *TAF6* or *TAF9* mRNAs. Consistent with our forward RIPs, no reciprocal mRNA enrichment was detected in any DUB pull-down (Fig 7H).

Together, these results demonstrate that while the DUB module relies on an obligate, sequential co-translational pathway for its own internal assembly, it does not assemble co-translationally onto the SAGA core. Instead, the DUB module incorporates into SAGA via a post-translational mechanism, establishing a distinct spatio-temporal boundary between intra-module biogenesis and holo-complex integration.

## Discussion

Our findings establish that the biogenesis of the yeast SAGA DUB module is governed by an obligate, linear, and unidirectional co-translational assembly pathway (Sus1 → Sgf11 → Ubp8 → Sgf73). Crucially, systematic genetic and in vivo interaction analyses demonstrate that when an upstream co-translational engagement is disrupted, downstream subunits fail to associate post-translationally. This stands in sharp contrast to in vitro biochemical reconstitutions, where recombinant Ubp8 and an N-terminal fragment of Sgf73 purified separately from Escherichia coli can associate post-translationally with a pre-formed Sus1–Sgf11 heterodimer to form a catalytically active tetramer (43,48), and where heterologous co-expression of these components in bacteria yields the intact core module (43,44). Together, our results suggest that within the crowded eukaryotic intracellular environment—characterized by separate monocistronic transcripts, low physiological subunit concentrations, competing interaction networks, and active orphan-protein quality control—spontaneous post-translational encounters are insufficient to assemble full-length DUB subunits into productive complexes. Moreover, sequential co-translational engagement may prevent kinetically trapped or non-native conformations that could otherwise arise during post-translational folding of full-length partners. Thus, for the DUB module, co-translational biogenesis does not merely enhance assembly kinetics but appears to act as an essential architectural determinant that dictates complex formation in vivo.

While the initial co-translational engagement between Sus1 and nascent Sgf11 is evolutionarily conserved in human cells via their orthologs ENY2 and ATXN7L3 (10), our data reveal an apparent difference further downstream. In *S. cerevisiae*, Sus1 also associates with translating *UBP8* and *SGF73* mRNAs, whereas ENY2 immunoprecipitates in human cells showed no enrichment of *USP22* mRNA and no clear enrichment of *ATXN7* mRNA (10). This difference could reflect technical factors – such as variations in translation kinetics or immunoprecipitation sensitivity – or alternatively represent genuine evolutionary specialization. Unlike yeast, humans express two additional ubiquitin proteases, USP27X and USP51 – encoded by X-linked retrogenes derived from a USP22 ancestor (52) – that compete with USP22 for ENY2 – ATXN7L3 binding to form SAGA-independent DUB complexes (38). Relaxing the co-translational coupling of ENY2-ATXN7L3 to nascent USP22 would therefore provide the plasticity required to assemble alternative deubiquitinase modules. Evolution may also have layered regulatory control directly onto the conserved first step of assembly: humans harbor a cytoplasmic paralog of ATXN7L3, ATXN7L3B, which competes with ATXN7L3 for ENY2 binding without interacting with USP22 or ATXN7, thereby limiting SAGA DUB assembly and modulating global H2Bub1 levels (53). Whether ENY2 binds ATXN7L3B co-or post-translationally remains unknown, but its cytoplasmic localization raises the possibility that ATXN7L3B sequesters ENY2 in the cytoplasm to directly throttle its co-translational recruitment to nascent ATXN7L3.

Within yeast, our hierarchical model (Sus1 → Sgf11 → Ubp8 → Sgf73) accounts for both the physical persistence and the functional maturation of DUB subcomplexes. While Co-IP assays showed that the Sus1-Sgf11-Ubp8 trimer remains physically intact in *sgf73*Δ soluble extracts ((46,47); Fig 5A,B,D), live-cell NanoBiT assays revealed opposing shifts in subunit complementation in the absence of Sgf73, decreasing the Sus1–Ubp8 interaction index while increasing that of Sgf11–Ubp8 (Fig 6B,D). Because RIP-qPCR positions Sgf73 as the terminal subunit recruited during translation, these divergent NanoBiT signals most likely reflect changes in the relative spatial proximity or orientation of the subunits – and consequently of their fused luciferase fragments – within the incomplete trimer. This interpretation aligns with a post-assembly role for Sgf73 in compacting and stabilizing the module into its catalytically competent conformation, consistent with crystal structures showing that Sgf73 contacts all three partners and bridges the assembly and catalytic lobes of Ubp8 (43,44,54).

This structural requirement also accounts for our steady-state protein abundance data. Sgf73 is the only subunit whose deletion reduces the abundance of an upstream partner, causing a moderate (∼30%) decrease in Ubp8 levels, likely due to partial destabilization of the catalytically inactive Sus1-Sgf11-Ubp8 trimer accumulated in *sgf73*Δ cells. By contrast, Sus1 and Sgf11 steady-state pools remain buffered against *SGF73* loss. Beyond potential sensitivity limits in detecting a minor drop in their abundance, biological partitioning likely protects these early subunits: Sus1 is a stable core component of the TREX-2 mRNA export complex (55), and Sgf11 genetically interacts with Sus1 in mRNA export independently of Ubp8 (46), pointing to shared or subcomplex-specific roles outside SAGA. Notably, proteasome-mediated remodeling of SAGA has been shown to liberate an active Sgf73-Ubp8 dimer (41), raising the possibility that a complementary Sus1-Sgf11 subcomplex also persists *in vivo* following module dissociation.

Conversely, upstream subunits exert a strong stabilizing influence on downstream partners – with the notable exception of Sgf73, whose steady-state abundance is entirely self-sustained – and our linear pathway unifies over fifteen years of biochemical and genetic observations on SAGA composition and function. Loss of *SUS1* severely compromises Sgf11 and reduces Ubp8 accumulation, whereas *SGF11* deletion diminishes Ubp8 levels, indicating that early co-translational engagements are required to sustain productive synthesis, proper folding, or protection from degradation. Furthermore, because downstream assembly cannot proceed without an intact upstream platform, our model directly explains why Ubp8-TAP purifications in *sgf11*Δ strains recover only isolated Ubp8, whereas Ubp8-TAP in *sgf73*Δ cells co-purifies the Sus1-Sgf11-Ubp8 trimer (47). Similarly, it accounts for why Sus1 purifications in *ubp8*Δ mutants still retain Sgf11 yet – as in *sgf11*Δ cells – fail to recover Sgf73 or core SAGA subunits, and why mass spectrometry of SAGA purified from *sus1*Δ or *ubp8*Δ cells detects Sgf73 anchored to the SAGA core in the complete absence of Sus1, Sgf11, and Ubp8 (46–48). By dictating whether the DUB module can bridge to Sgf73 and dock onto SAGA, this assembly hierarchy also provides a mechanistic basis for the codependent recruitment of Sus1 and Ubp8 to target promoters such as GAL1 and the dependence of proper Sus1 intranuclear localization on Sgf11 and Ubp8 (46).

A central finding of this work is the mechanistic boundary separating intra-module DUB biogenesis from holo-SAGA integration. Despite extensive structural contacts between Sgf73 and the SAGA core interface formed by Spt20, Ada1, Taf12, Taf6, and Taf9 (31), we detected no co-translational interactions between core and DUB subunits, strongly indicating that the DUB module docks onto SAGA post-translationally. This contrasts with the HAT module, where Ada2 engages nascent Spt20 co-translationally (35), coupling HAT biogenesis directly to SAGA core assembly. Instead, this post-translational docking mode parallels the biogenesis of Tra1, which undergoes co-translational folding assisted by the Hsp90 co-chaperone TTT before post-translational incorporation into SAGA or NuA4 (50,56). Thus, SAGA integrates distinct assembly strategies – co-translational subunit templating (DUB), direct co-translational core tethering (HAT), and chaperone-assisted folding prior to docking (Tra1) – within a single macromolecular machine. Moreover, assembling the DUB module as an autonomous, SAGA-independent unit (Fig 8) provides a direct biosynthetic route to generate free, functional DUB complexes capable of operating outside SAGA (39–41,57–59).

**Figure 8.**
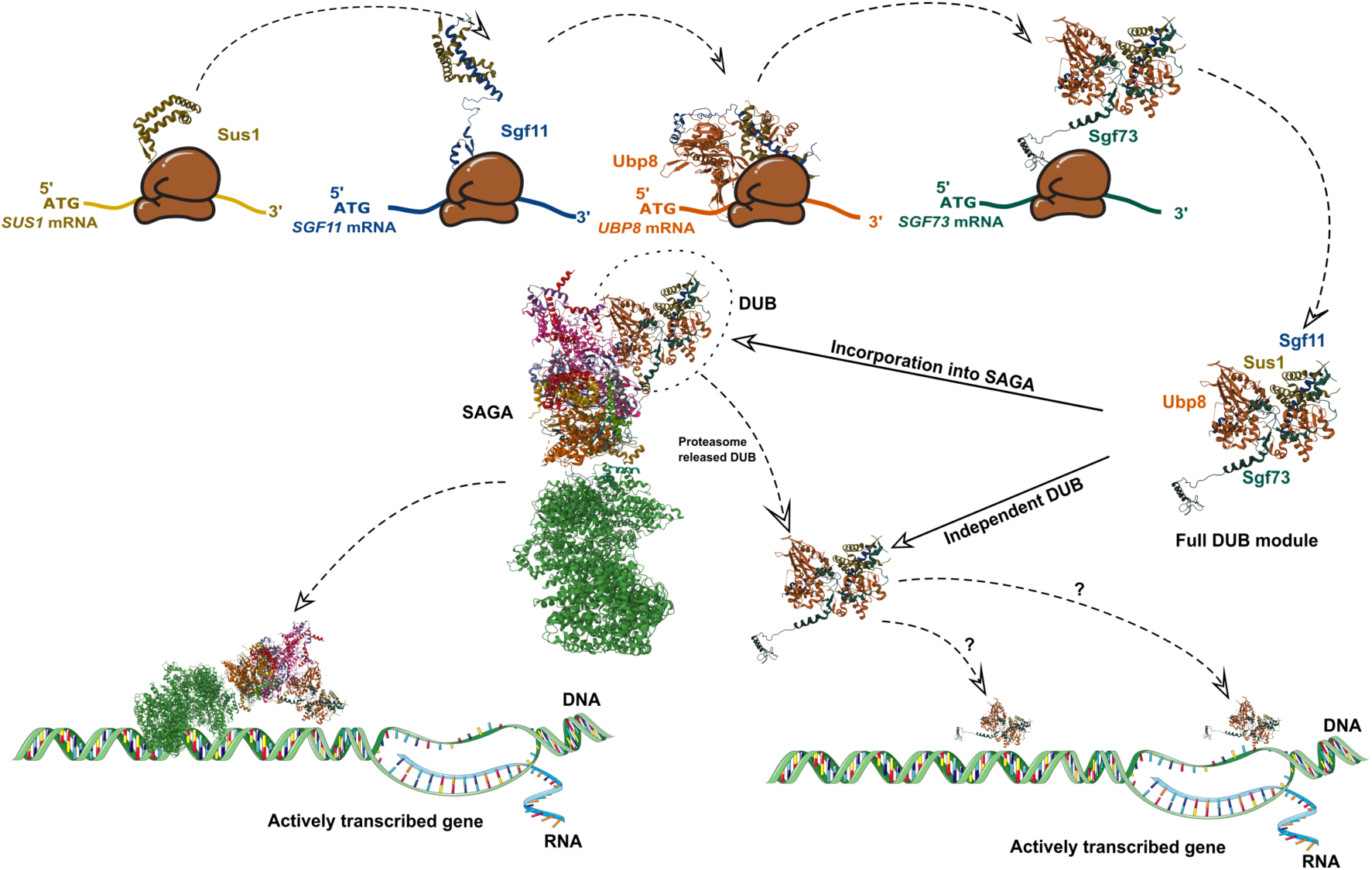
Working model for autonomous co-translational assembly of the yeast SAGA DUB module and its partitioning into SAGA-associated and SAGA-independent pools. Top: Sequential co-translational assembly of the four-subunit DUB module occurs independently of the SAGA core complex. Fully synthesized Sus1 co-translationally engages nascent Sgf11 emerging from the translating ribosome; the resulting Sus1–Sgf11 heterodimer subsequently binds nascent Ubp8, and the pre-assembled Sus1–Sgf11–Ubp8 trimer co-translationally recruits nascent Sgf73 to generate a complete, autonomous DUB module (Full DUB module). Structural representations of the individual DUB subunits and the holo-SAGA complex are derived from the *Saccharomyces cerevisiae* SAGA cryo-EM structure (PDB: 6T9I). Bottom: Following assembly, the intact DUB module can dock onto the SAGA core (Incorporation into SAGA) to act as part of the holo-SAGA complex on actively transcribed genes (bottom left). In addition, we propose that a pool of newly assembled DUB modules may remain unattached to SAGA (Independent DUB), converging with DUB modules dissociated from SAGA via proteasome-mediated remodeling (Proteasome released DUB) to carry out putative SAGA-independent roles at actively transcribed chromatin (bottom right; dashed arrows labeled with question marks indicate hypothetical pathways).

Placed in a broader context, our findings expand the repertoire of co-translational assembly strategies described for transcriptional and chromatin-associated complexes (6,7,10,36,60) . In Set1C/COMPASS, multiple subunits converge directly onto nascent Set1, which serves as a central assembly scaffold (6). In TFIID, discrete structural building blocks—formed through mixed pairwise co- and post-translational interactions—dock co-translationally onto emerging domains of the central scaffold TAF1 (60). The SAGA DUB module illustrates a distinct modular logic: rather than relying on a single central nascent scaffold or docking co-translationally onto a core subunit akin to TAF1, an entire autonomous catalytic module is assembled through a linear four-step co-translational relay prior to post-translational integration into the SAGA core. Together, these architectures highlight how eukaryotic cells tailor co- and post-translational assembly routes to balance modular autonomy with holo-complex fidelity.

Altogether, our work illustrates how a multi-step, unidirectional co-translational cascade governs the biogenesis of an entire functional module in vivo, channeling nascent subunits toward productive assembly while establishing the architectural prerequisites for subsequent post-translational docking onto a megadalton complex. Defining the nascent-chain determinants, folding intermediates, and ribosome-proximal factors that license each sequential engagement will be an important next step toward understanding how translational dynamics shape quaternary structure. More broadly, comparing assembly pathways across evolutionarily divergent eukaryotes will reveal how variations in subunit domain architecture, paralog diversification, and gene expression regimes rewire the balance between co- and post-translational assembly.

## Material and methods

### Yeast growth conditions

Standard culture media and genetic manipulations were used. *S. cerevisiae* strains were grown in YPAD (1% yeast extract, 1% adenine, 2% peptone and 2% glucose; supplemented with 2% agar for solid media) at 30 °C over-night, diluted and grown to mid-log phase (∼1 × 10^7^ cells per ml). For serial dilution drop tests, exponentially growing cultures were adjusted to OD_600_ = 1. Then, 5 µl of 1, 10, 10^2^, 10^3^ and 10^4^ dilutions were spotted on YPAD plates (control) or YPAD plates containing 200 mM HU or 3% formamide. Cells were grown for at least 48 h at 30 °C before imaging.

### Construction of yeast strains

*S. cerevisiae* strains used are listed in Supplementary Table S1 and were constructed by standard procedures, using either chemical transformation or genetic crosses. Strains with gene deletions or C-terminally epitope-tagged proteins were constructed by PCR-based gene targeting of the respective open-reading frame (ORF) with kanMX6, natMX6 or hphMX6 cassettes, amplified from pFA6a backbone plasmids (see below and (61,62)). DNA fragments used for homologous recombination were generated by PCR, Gibson assembly cloning (kit E2611L, New England Biolabs), or gene synthesis. Cloning strategies and primers were designed using the sequences available at Saccharomyces Genome Database (SGD) (63). All relevant primer sequences are listed in Supplementary Table 1. Transformants were screened for correct integration by PCR and, when appropriate, verified by Sanger sequencing or Western blotting. For each transformation, 2–6 individual clones were purified and analyzed.

### Plasmid construction

C-terminal tagged strains used for RIP assays contained a -GS-HA3-TAP2 tag derived from pEV58, which we constructed from a pFA6a backbone plasmid (50,56) by inserting a 4 glycines followed by 4 serines (GGGGSSSS) linker in fusion with the -HA3-TAP2 epitope using a synthetized linker and the Gibson assembly kit (E2611L, New England Biolabs). FLAG tagged strains contained a 6 glycines linker followed by 3 FLAG epitope derived from pFA6a-6xGLY– 3xFLAG-hphMX4 plasmid (64). Alternatively, MYC tagged strains were constructed using a pFA6a-13xMYC-natMX6 plasmid (Euroscarf).

### Protein extraction

Protein extracts were prepared from 1 or 50 ml of exponentially growing cells, using either trichloroacetic acid (TCA) precipitation or standard lysis buffer (LB: 50 mM Tris-HCl pH 7.5, 100 mM NaCl, 1.5 mM MgCl_2_ and 0.15% NP-40), respectively. For standard lysis, LB was supplemented with protease inhibitors, including cOmplete EDTA-free cocktails tablets (04693132001, Roche), 100 mM PMSF, 1 mg/ml bestatin, and 1 mg/ml pepstatin A, as well as phosphatase inhibitors, including 50 mM β-glycerolphosphate, 1 mM NaF, and 1 mM Na_3_VO_4_. Glass beads were added to the cell suspension, and lysis was performed using FastPrep (MP Biomedicals) in 4 cycles of 30 seconds at 5.5 power with 1 minute rest on ice in between. Beads were removed by centrifugation for 1 minute a 1000 rpm, 4°C, and extracts were cleared by centrifugation at 14000 rpm, 4°C, for 10 minutes. For TCA extraction, cells were lysed adding 1.85 M NaOH, vortex and kept in ice for 10 minutes. Then, 55% TCA was added, mixed by microtube inversion, followed by an incubation in ice for 10 minutes. Protein extract was pelleted by centrifugation at 14000 rpm, 4°C, for 15 minutes. Protein pellet was resuspended in HU-buffer containing 8 M urea, 20% SDS, 200 mM Tris pH 6.8, 1mM EDTA pH 8, 1.5% DTT and 0.1% bromophenol-blue. Resuspension volume was adjusted depending on the initial OD of the culture for normalization. Samples were boiled for 10 minutes at 95 °C. 5 μl of each sample was loaded on a 10% SDS–acrylamide gel. Stain-free gels were used to normalize for protein amounts across samples.

### Western blot

Western blotting was performed using a anti-HA antibody for detection of the HA tag (16B12, BioLegend), an anti-FLAG antibody for detection of the FLAG tag (M2, Sigma), an anti-G6PDH antibody for detection of endogenous *S. cerevisiae* G6PDH or an anti-Pgk1 antibody for detection of endogenous Pgk1 protein. Protein concentrations were measured by the Bradford method and used to load equal amounts of proteins across samples. G6PDH, Pgk1 and Ponceau red were used as loading controls. Visualization was performed using a ChemiDoc system (BioRad). Quantification of signal intensity was performed using non-saturated exposures and analyzed using Fiji (65).

### Co-immunoprecipitation

For co-immunoprecipitation assays, proteins were extracted using LB lysis buffer as indicated above and quantified using the Bradford method. For immunoprecipitation (IP) 2-4 mg of total protein extract were incubated with 1 µg of anti-FLAG, anti-HA or anti-MYC antibodies for 1 h at room temperature (RT). Samples were then mixed with Protein G magnetic dynabeads (Invitrogen) and incubated for an additional 1 h at RT. Beads were then washed three times with lysis buffer prior to elution in 30 µl of Laemmli buffer and boiling 10 minutes at 95 °C. Proteins were visualized by Western blotting as previously described. Input samples corresponded to 1.25-2.5% (25-50 µg) of the volume used for IP.

### RT-qPCR analyses

Reverse transcription and quantitative PCR analyses of cDNA were performed using RNA extracted from 50 mL of exponentially growing cells using the Invitrogen Purelink RNA Mini Scale kit. DNA was removed by DNase I digestion, using the TURBO DNA-free kit on 10 µg of total RNA. 1 μg of DNase-treated RNA was then reverse transcribed (RT) at 55°C with random hexanucleotide primers (Roche), using the SuperScript III First-Strand System. Fluorescence-based quantitative PCR was performed with SYBR Green and used to calculate relative cDNA quantities, from the slope produced by standard curves for each primer pair, in each experiment. DNase-treated RNA samples were used as controls for the presence of genomic DNA contaminants. Standard curve slopes were comprised between −3.5 (90% efficiency) and −3.15 (110% efficiency), with an r^2^ > 0.9. Primer sequences are listed in Supplementary Table S2.

### RNA-IP (RIP)

RIP experiments were done as described previously (7,66), with minor modifications. Where indicated, exponentially growing cultures were treated with 1 mM for 15 min at 30 °C prior to harvesting to dissociate translating ribosomes. Briefly, 50 mL of exponentially growing cells were harvested, washed with 5 ml of ice-cold water, resuspended in 200 µl of IPC buffer (20 mM Tris-HCl pH 8, 140 mM KCl, 1.8 mM MgCl_2_, 0,1 % NP-40, 0,2 mg/mL heparine) supplemented with protease and phosphatase inhibitors as indicated for protein extraction, DNase (2 µl of Turbo DNase kit, Invitrogen) and 1 µl of RNasin RNase inhibitor (Promega), and broken using a FastPrep (MP Biomedicals) as indicated for protein extraction. Immunoprecipitation (IP) was performed using 50 µl of Dynabeads Pan-Mouse IgG slurry, washed three times with IPC buffer, pre-incubated with 1 μg of anti-HA (16B12) antibody overnight at 4°C and washed again the same way before adding the cell extract. 2-4 mg of total protein extracts were used in each IP for 2 hours at 4°C. Input (IN) and IP RNAs were extracted using the Invitrogen Purelink RNA Mini and Micro Scale kits, respectively. DNase treatment, reverse transcription, and qPCR analyses of cDNA were performed as described for RT-qPCR analysis. IN samples were diluted 50-fold while IP samples were diluted 2-to-5-fold. Fold enrichment over no tag was calculated by the DDCt method, correcting input and IP Ct values for their corresponding dilution factor. DCt was calculated by substracting the input Ct value from the IP Ct value of each sample. DDCt was calculated by substracting the mean DCt value from non-tagged samples (non-paired samples) from each DCt value of each sample. To determine the specificity of enrichment of the tagged protein, the corresponding untagged control samples were included in each RIP experiment. mRNAs corresponding to bait proteins were used as positive controls for RIP performance. All primer sequences are listed in Supplementary Table S1.

### NanoLuc luminescence measurements

To perform NanoLuc luminescence measurements, cells were cultured in YPAD. From exponentially growing cell culture, OD600 was measured and adjusted to 0.8 OD in 1 mL YPAD. Next, 300 uL of the adjusted culture was transferred to a translucent 96-well plate for OD measurement. A Promega GloMax luminometer was used to record OD_600_ with a previous shake of 30 s at 500 cycles per minute. Then, 100 uL of the cell culture from the translucent 96-well plate was transferred to an opaque white 96-well plate for luminescence measurement. Each cell culture was assayed in duplicated wells: one 100 uL in a well was destinated to furimazine addition, while the other 100 uL of cells in a different well, served as a control without furimazine. Each 100 uL of cells were permeabilized by adding 25 μL of Passive Lysis Buffer (Promega) and incubated at room temperature for 5 minutes. Following incubation, 125 uL of furimazine solution (1:50 dilution of a 5 mM furimazine stock in TBST-Thiourea buffer, obtaining a 0.04 mM furimazine as final concentration per well) or alternatively 125 uL of control solution (1:50 dilution of 85% ethanol – 15% glycerol mixture in TBST-Thiourea buffer) without furimazine, was added to the respective wells. Luminescence was then measured using the Promega GloMax luminomener with 1 s integration time, preceded by a 30 s shake at 500 cycles per minute before luminescence measurement. The luminescence signal was normalized against the OD measurement of each well and corrected for the background luminescence signal of a non-tagged NanoLuc strain.

### NanoBiT assays

Similarly to NanoLuc luminescence measurements, for NanoBiT assays cells were cultured in YPAD. From exponentially growing cell culture, OD_600_ was measured and adjusted to 0.8 OD in 1 mL YPAD. Next, 300 uL of the adjusted culture was transferred to a translucent 96-well plate for OD measurement as described previously. Similarly, 100 uL of cells were then transferred to an opaque white 96-well plate for luminescence measurement in duplicate wells for furimazine and control solutions, respectively. Subsequently, 150 uL of furimazine solution (1:100 dilution of 5 mM furimazine stock in YPAD, obtaining a 0.04 mM furimazine as final concentration per well) or 150 uL of control solution (1:100 dilution of 85% ethanol – 15% glycerol mixture in YPAD) was added to intact non-permeabilized cells to the respective wells. Luminescence was then measured using the Promega GloMax luminometer as described previously for NanoLuc luminescence measurements. The luminescence signal was normalized against the OD_600_ measurement of each well, and background subtraction using the corresponding single-LgBiT tagged control strain was applied for calculating the normalized interacting index (Nii, see Statistical analyses).

### Statistical analyses

Statistical analyses were performed using GraphPad Prism. For RIP-qPCR assays, relative fold enrichment values (2^-ΔΔCt^) were log_2_-transformed and analyzed by two-way ANOVA (factors: transcript and strain/treatment) followed by Tukey’s multiple comparisons test assessing simple effects within each transcript. Statistical comparisons were omitted in panels where target mRNA enrichment remained at background levels comparable to the negative control *VTC3*, or where sample size was *n* < 3. For NanoLuc luminescence assays, normalized RLU/OD_600_ values (expressed relative to the wild-type control) were analyzed by one-way ANOVA followed by Dunnett’s multiple comparisons test against wild type. For NanoBiT assays, the limit of detection (LoD) was defined as the mean + 3 SD of the corresponding single-subunit LgBiT-only negative control (lacking a SmBiT partner). Raw luminescence values normalized to culture density (RLU/OD_600_) were log^10^-transformed and analyzed by one-way ANOVA followed by Šídák’s multiple comparisons test. To account for differences in subunit steady-state abundance across deletion backgrounds, a normalized interacting index (Nii) was calculated for each biological replicate by: (i) subtracting the mean RLU/OD_600_ of the single-LgBiT negative control, (ii) normalizing background-subtracted values to the wild-type strain within the same biological replicate (WT = 1), and (iii) dividing the resulting relative interaction value by the product of the normalized steady-state abundance of both interacting subunits in that mutant background (as independently determined by NanoLuc assays). Untransformed Nii values were evaluated using two-tailed one-sample *t*-tests against theoretical means of 1.0 (WT) and LoD. Due to the high sensitivity and wide dynamic range of RIP-qPCR and luminescence-based assays, a stringent significance threshold of α = 0.01 was applied. Western blot densitometry and RT-qPCR data were analyzed using two-tailed unpaired Welch’s *t*-tests with a significance threshold of α = 0.05. Sample sizes (*n*, representing independent biological replicates) and exact *p*-values are reported in the figure legends.

## Acknowledgments

We thank Dominique Helmlinger for critical reading of the original manuscript draft and for sharing plasmids used alongside this study. We are grateful to Gwenaël Rabut for supplying NanoLuc and NanoBiT strains and plasmids and for advice on their implementation. We thank members of laboratories 201 and 212 at IBiS for helpful discussions. This work is supported by the EMERGIA grant number EMC21_00283, from the Consejería de Universidad, Investigación e Innovación of the regional government Junta de Andalucía, to AEV, and by in-house funding of the University of Seville through the Programa al Estímulo de Áreas con Necesidades Investigadoras y de la Actividad Investigadora Emergente of the VII Plan Propio de Investigación y Transferencia de la Universidad de Sevilla (PPIT-US), calls 2023 and 2025, to AEV. ABM was supported by a contract associated to the EMERGIA grant EMC21_00283. JAHS was supported through a FPU fellowship (FPU24/04574) from the Ministerio de Ciencia, Innovación y Universidades. LMA was supported through a Beca de Colaboración from the Ministerio de Educación, Formación Profesional y Deportes. We thank all funders for their contribution.

## Author contributions

Conceptualization: AEV. Data curation: ABM, JAHS, AEV. Formal analysis: ABM, AEV. Funding acquisition: AEV, JAHS, LMA. Investigation: ABM, JAHS, LMA. Methodology: ABM, JAHS, LMA, AEV. Project administration: AEV. Supervision: AEV. Validation: ABM, JAHS, LMA. Visualization: AEV, ABM. Writing – original draft: AEV. Writing – review and editing: All authors.

