## Supplementary material for "Sequential co-translational assembly of the yeast SAGA DUB module precedes post-translational core docking"

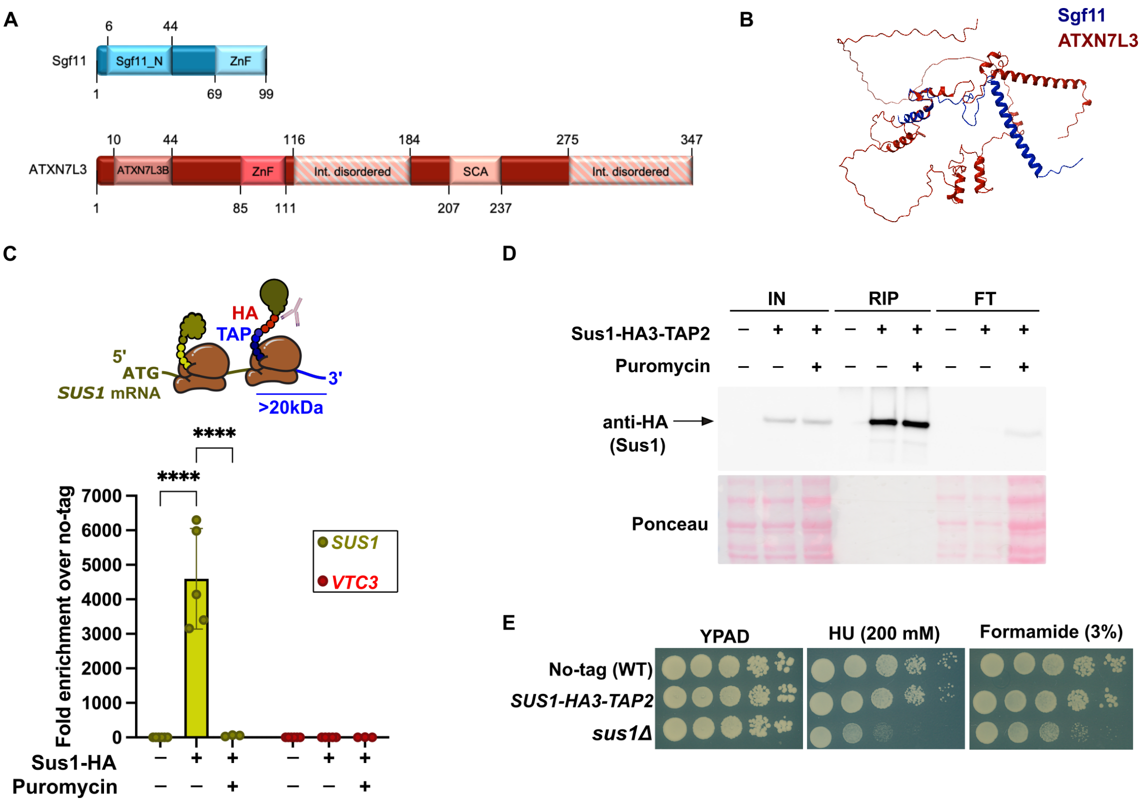


**Supplementary Figure S1. Structural comparison between yeast Sgf11 and human ATXN7L3, and functional validation of the C-terminal HA3-TAP2 tagging strategy. (A)** Schematic domain organization of *Saccharomyces cerevisiae* Sgf11 (99 amino acids) and *Homo sapiens* ATXN7L3 (347 amino acids), depicting the conserved N-terminal (Sgf11_N / ATXN7L3B) and zinc-finger (ZnF) domains alongside human-specific intrinsically disordered regions and the SCA region. Amino acid boundary positions are indicated. **(B)** Structural superposition of yeast Sgf11 (blue) and human ATXN7L3 (dark red), illustrating structural conservation across their core folded domains. **(C)** *Top:* Schematic illustrating co-translational self-immunoprecipitation of *SUS1* mRNA via the C-terminal HA3-TAP2 tag (>20 kDa), which emerges from the ribosomal exit tunnel once the Sus1 polypeptide is fully synthesized. *Bottom:* Anti-HA RIP-qPCR analysis of *SUS1* mRNA enrichment alongside the unrelated negative control *VTC3* in untagged (−), Sus1-HA3-TAP2 (+), and puromycin-treated Sus1-HA3-TAP2 (+ Puromycin) strains. Individual data points represent independent biological replicates ($n$ = 5), with mean ± SD overlaid, expressed as fold enrichment ($2^{-\Delta\Delta Ct}$) relative to the mean $\Delta Ct$ of the untagged control. Individual replicates of this control are plotted to illustrate baseline variability. Statistical comparisons shown for *SUS1* (untagged vs. Sus1-HA, and Sus1-HA vs. Sus1-HA + puromycin) were determined by two-way ANOVA on $\log_2$-transformed relative enrichment values followed by Tukey's multiple comparisons test (simple effects within each transcript), using a significance threshold of $\alpha = 0.01$ (****$p < 0.0001$); *VTC3* is plotted alongside as an unrelated negative control to illustrate baseline background. **(D)** Western blot control of the anti-HA RIP assay in (C), analyzing input (IN), immunoprecipitated (RIP), and flow-through (FT) fractions from untagged (−), Sus1-HA3-TAP2 (+), and puromycin-treated Sus1-HA3-TAP2 (+ Puromycin) cell extracts. Membranes were probed with anti-HA to detect Sus1-HA3-TAP2 and stained with Ponceau S to verify protein loading and transfer. **(E)** Serial dilution spot assays assessing cell growth of untagged wild-type (No-tag WT), *SUS1-HA3-TAP2*, and *sus1Δ* strains on rich medium (YPAD) and under genotoxic or transcriptional stress conditions (200 mM hydroxyurea [HU] or 3% formamide), confirming that the C-terminal HA3-TAP2 tag preserves Sus1 function *in vivo*.

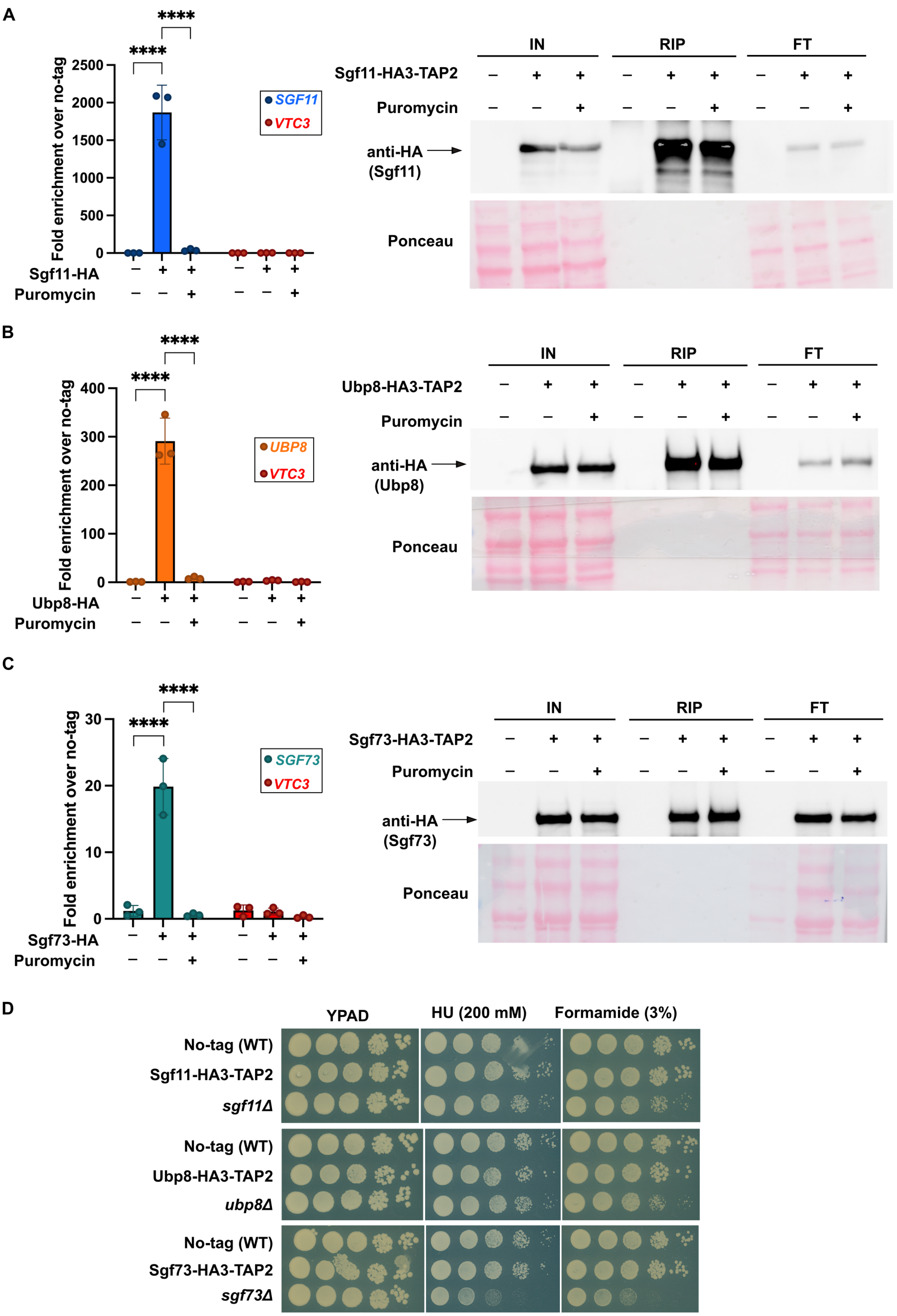


**Supplementary Figure S2. Co-translational self-mRNA association and functional validation of C-terminally HA3-TAP2-tagged Sgf11, Ubp8, and Sgf73. (A–C)** Co-translational self-immunoprecipitation and Western blot validation of Sgf11-HA3-TAP2 (A), Ubp8-HA3-TAP2 (B), and Sgf73-HA3-TAP2 (C). *Left:* Anti-HA RIP-qPCR analysis of *SGF11* (A), *UBP8* (B), and *SGF73* (C) mRNA enrichment alongside the unrelated negative control *VTC3* in untagged (−), HA3-TAP2-tagged (+), and puromycin-treated HA3-TAP2-tagged (+ Puromycin) strains. Individual data points represent independent biological replicates (*n* = 3), with mean ± SD overlaid, expressed as fold enrichment ($2^{-\Delta\Delta Ct}$) relative to the mean $\Delta Ct$ of the untagged control. Individual replicates of this control are plotted to illustrate baseline variability. Statistical comparisons shown for the target DUB transcripts (untagged vs. tagged, and tagged vs. tagged + puromycin) were determined by two-way ANOVA on $\log_2$-transformed relative enrichment values followed by Tukey's multiple comparisons test (simple effects within each transcript), using a significance threshold of $\alpha = 0.01$ (****$p < 0.0001$); *VTC3* is plotted alongside as an unrelated negative control to illustrate baseline background. *Right:* Western blot controls of the corresponding anti-HA RIP assays analyzing input (IN), immunoprecipitated (RIP), and flow-through (FT) fractions, probed with anti-HA and stained with Ponceau S to verify protein loading and transfer. **(D)** Serial dilution spot assays assessing cell growth of untagged wild-type (No-tag WT), HA3-TAP2-tagged (Sgf11-HA3-TAP2, Ubp8-HA3-TAP2, Sgf73-HA3-TAP2), and deletion mutant (*sgf11*Δ, *ubp8*Δ, *sgf73*Δ) strains on rich medium (YPAD) and under genotoxic or transcriptional stress conditions (200 mM hydroxyurea [HU] or 3% formamide), confirming that C-terminal HA3-TAP2 tagging preserves subunit function *in vivo*.

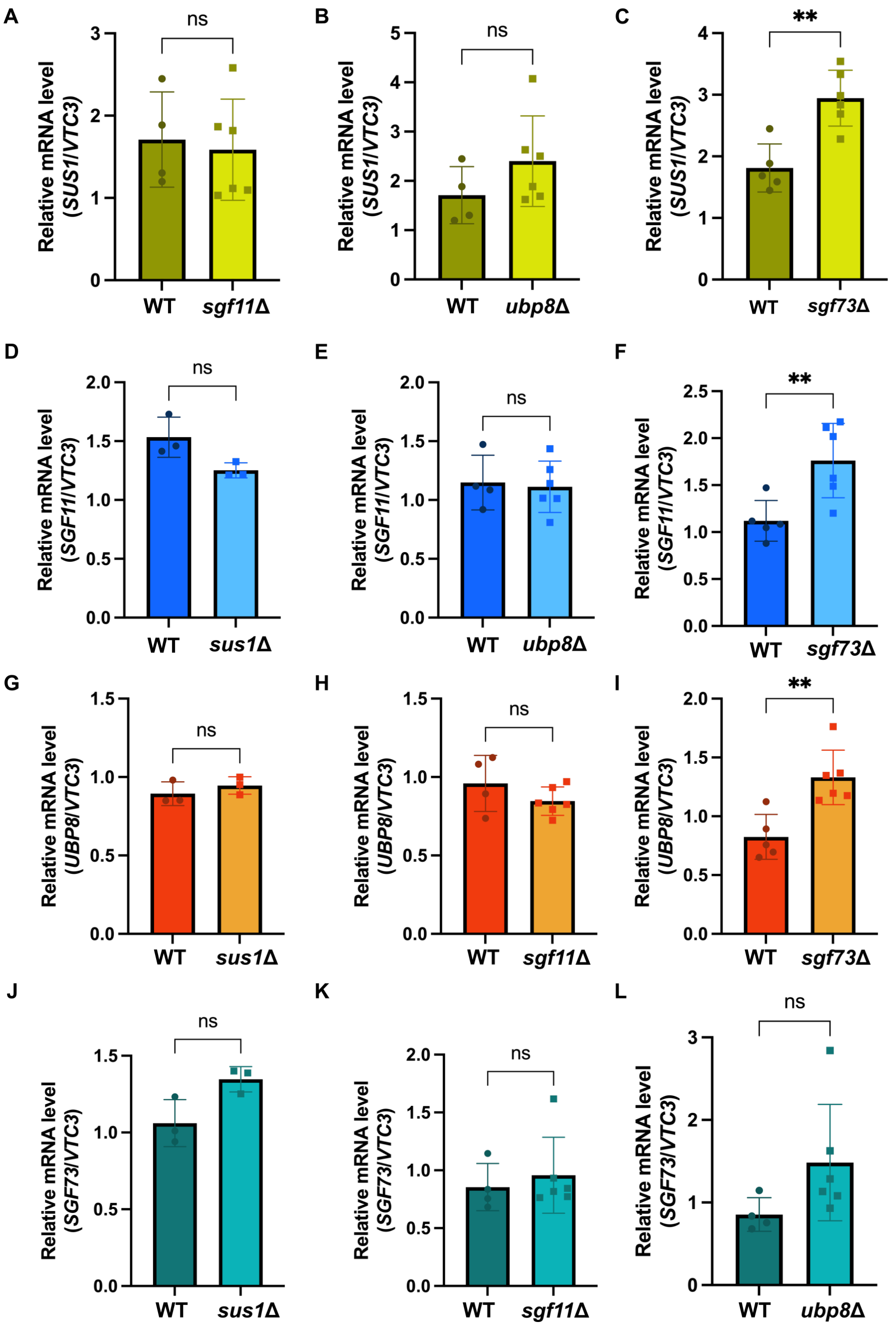


**Supplementary Figure S3. Steady-state mRNA levels of DUB module subunits in single deletion mutants confirm post-transcriptional regulation of protein abundance. (A–L)** RT-qPCR analysis of relative steady-state mRNA levels of *SUS1* (A–C), *SGF11* (D–F), *UBP8* (G–I), and *SGF73* (J–L) in wild-type (WT) *versus* the indicated deletion mutant strains within the corresponding -HA3-TAP2-tagged backgrounds (matching Figure 3A–L): *SUS1* mRNA in Sus1-HA3-TAP2 WT *versus sgf11*Δ (A; $p = 0.7562$), *ubp8Δ* (B; $p = 0.1830$), and *sgf73*Δ (C; $p = 0.0016$); *SGF11* mRNA in Sgf11-HA3-TAP2 WT *versus sus1*Δ (D; $p = 0.0894$), *ubp8*Δ (E; $p = 0.8108$), and *sgf73*Δ (F; $p = 0.0094$); *UBP8* mRNA in Ubp8-HA3-TAP2 WT versus *sus1*Δ (G; $p = 0.3938$), *sgf11*Δ (H; $p = 0.3093$), and *sgf73*Δ (I; $p = 0.0032$); and *SGF73* mRNA in Sgf73-HA3-TAP2 WT versus *sus1*Δ (J; $p = 0.0630$), *sgf11*Δ (K; $p = 0.5590$), and *ubp8*Δ (L; $p = 0.0842$). Relative mRNA levels were quantified from qPCR standard-curve-derived quantities and normalized to *VTC3* as an internal reference control ("*GENE*"/*VTC3*). Individual data points represent independent biological replicates ($n$ = 4-6 in A, B, E, H, K, L; $n$ = 5-6 in C, F, I; $n$ = 3 in D, G, J), with mean ± SD overlaid. Statistical significance was determined by a two-tailed unpaired Welch's $t$-test ($^{**}p < 0.01$; ns, not significant).

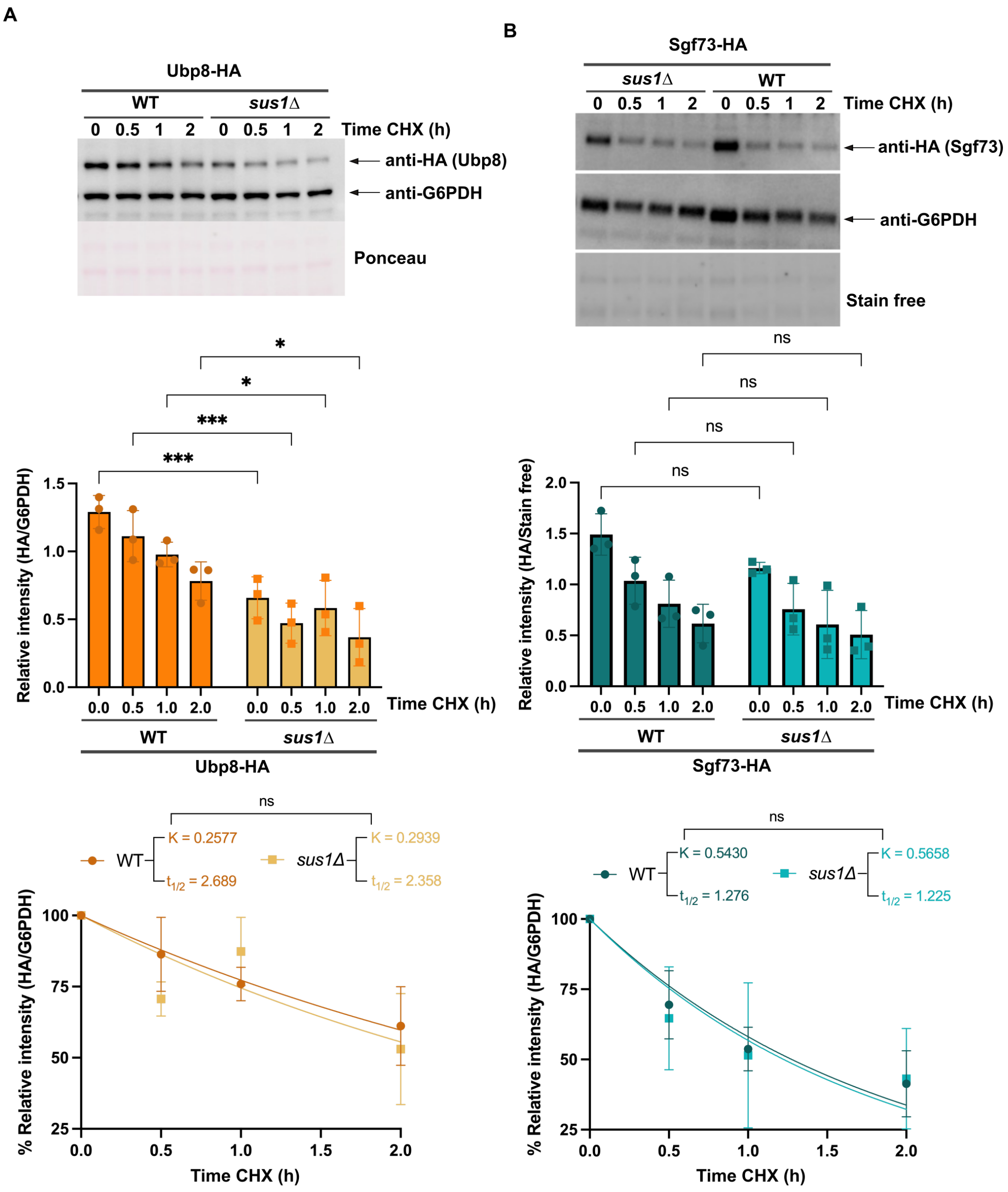


**Supplementary Figure S4. Cycloheximide chase analysis of Ubp8 and Sgf73 protein stability in wild-type and *sus1Δ* cells. (A, B)** Cycloheximide (CHX) chase assays monitoring the stability of Ubp8-HA3-TAP2 (Ubp8-HA; A) and Sgf73-HA3-TAP2 (Sgf73-HA; B) in wild-type (WT) and *sus1Δ* strains. *Top:* Representative Western blots of whole-cell extracts harvested at 0, 0.5, 1, and 2 h after CHX addition. Membranes were probed with anti-HA and anti-G6PDH (loading control) antibodies alongside Ponceau S staining. *Middle:* Densitometric quantification of Ubp8-HA (A) and Sgf73-HA (B) levels expressed as the unnormalized ratio of HA to G6PDH signal intensity (Relative intensity, HA/G6PDH). Individual data points represent independent biological replicates ($n$ = 3), with mean ± SD overlaid. Statistical significance between WT and *sus1*Δ at each time point was determined by two-way ANOVA followed by Šídák's multiple comparisons test (*$p < 0.05$, ***$p < 0.001$; ns, not significant; exact $p$ values for Ubp8-HA (A): 0 h, $p = 0.0008$; 0.5 h, $p = 0.0007$; 1 h, $p = 0.0353$; 2 h, $p = 0.0256$; exact $p$ values for Sgf73-HA (B): 0 h, $p = 0.3371$; 0.5 h, $p = 0.4915$; 1 h, $p = 0.7495$; 2 h, $p = 0.9656$). *Bottom:* Nonlinear regression analysis of protein decay kinetics. For curve fitting, values within each replicate were normalized to the corresponding $t = 0$ h time point (set to 100%) and fitted to a one-phase exponential decay model (constrained to $Y0 = 100$ and Plateau = 0). Calculated protein half-lives ($t_{1/2}$) and decay rate constants ($K$) for WT and *sus1Δ* are indicated on each graph.

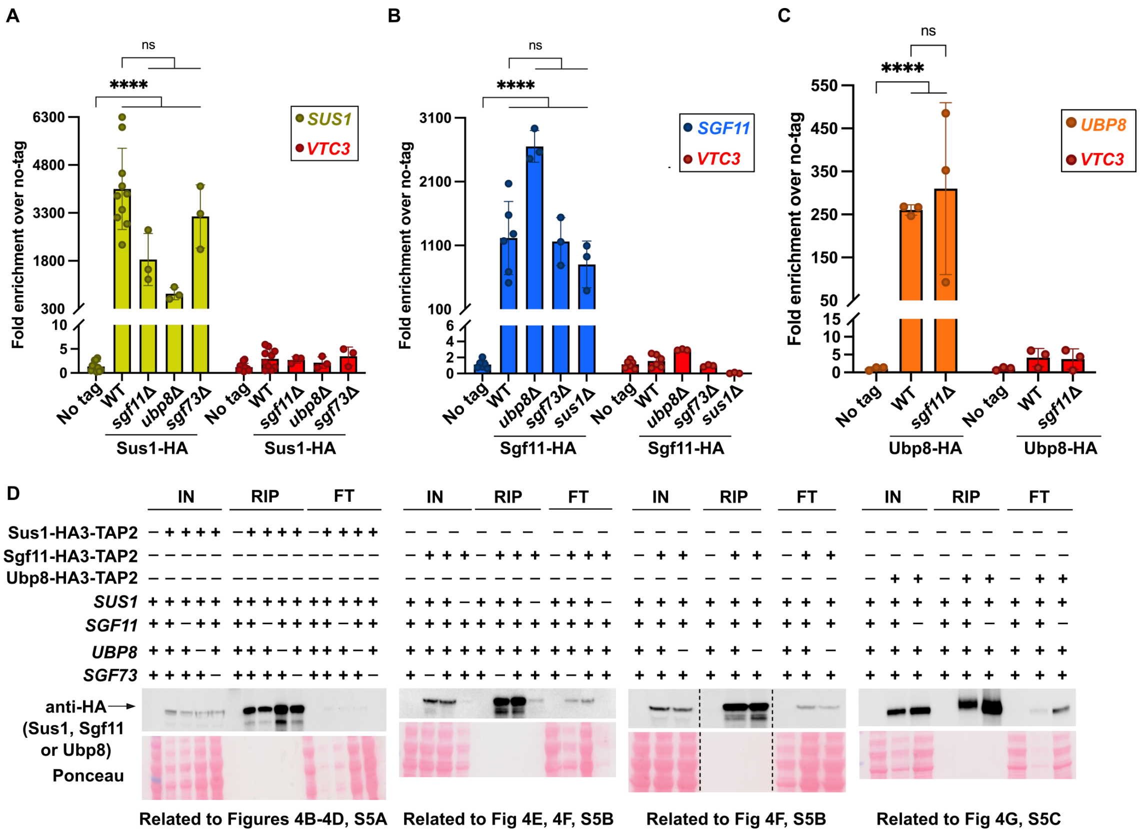


**Supplementary Figure S5. Co-translational self-mRNA association and immunoprecipitation controls of HA-tagged DUB subunits in deletion mutant backgrounds. (A–C)** Anti-HA RIP-qPCR control analysis quantifying co-translational self-mRNA enrichment alongside the unrelated negative control *VTC3* in untagged (No tag), wild-type (WT), and the indicated deletion mutant backgrounds: *SUS1* mRNA enrichment in Sus1-HA3-TAP2 immunoprecipitations (A), *SGF11* mRNA enrichment in Sgf11-HA3-TAP2 immunoprecipitations (B), and *UBP8* mRNA enrichment in Ubp8-HA3-TAP2 immunoprecipitations (C). Individual data points represent independent biological replicates (*n* = 3–9), with mean ± SD overlaid, expressed as fold enrichment ($2^{-\Delta\Delta Ct}$) relative to the mean $\Delta Ct$ of the untagged control. Individual replicates of this control are plotted to illustrate baseline variability. Statistical comparisons shown for the target DUB transcripts were determined across experimental conditions by two-way ANOVA on $\log_2$-transformed relative enrichment values followed by Tukey's multiple comparisons test (simple effects within each transcript), using a significance threshold of $\alpha = 0.01$ (*****p* < 0.0001; *ns*, not significant, $p \geq 0.01$); *VTC3* is plotted alongside as an unrelated negative control to illustrate baseline background. **(D)** Western blot controls for the anti-HA RIP-qPCR assays shown in Figure 4B–G and Supplementary Figure S5A–C, analyzing input (IN), immunoprecipitated (RIP), and flow-through (FT) fractions from untagged (−), wild-type (+), and the indicated DUB subunit deletion (−) strains. Membranes were probed with anti-HA to verify expression and immunoprecipitation efficiency of Sus1-HA3-TAP2, Sgf11-HA3-TAP2, and Ubp8-HA3-TAP2, and stained with Ponceau S to confirm protein loading and transfer. Vertical dashed lines in the third blot (related to Figs. 4F and S5B) indicate where intervening lanes from the same membrane were spliced out.

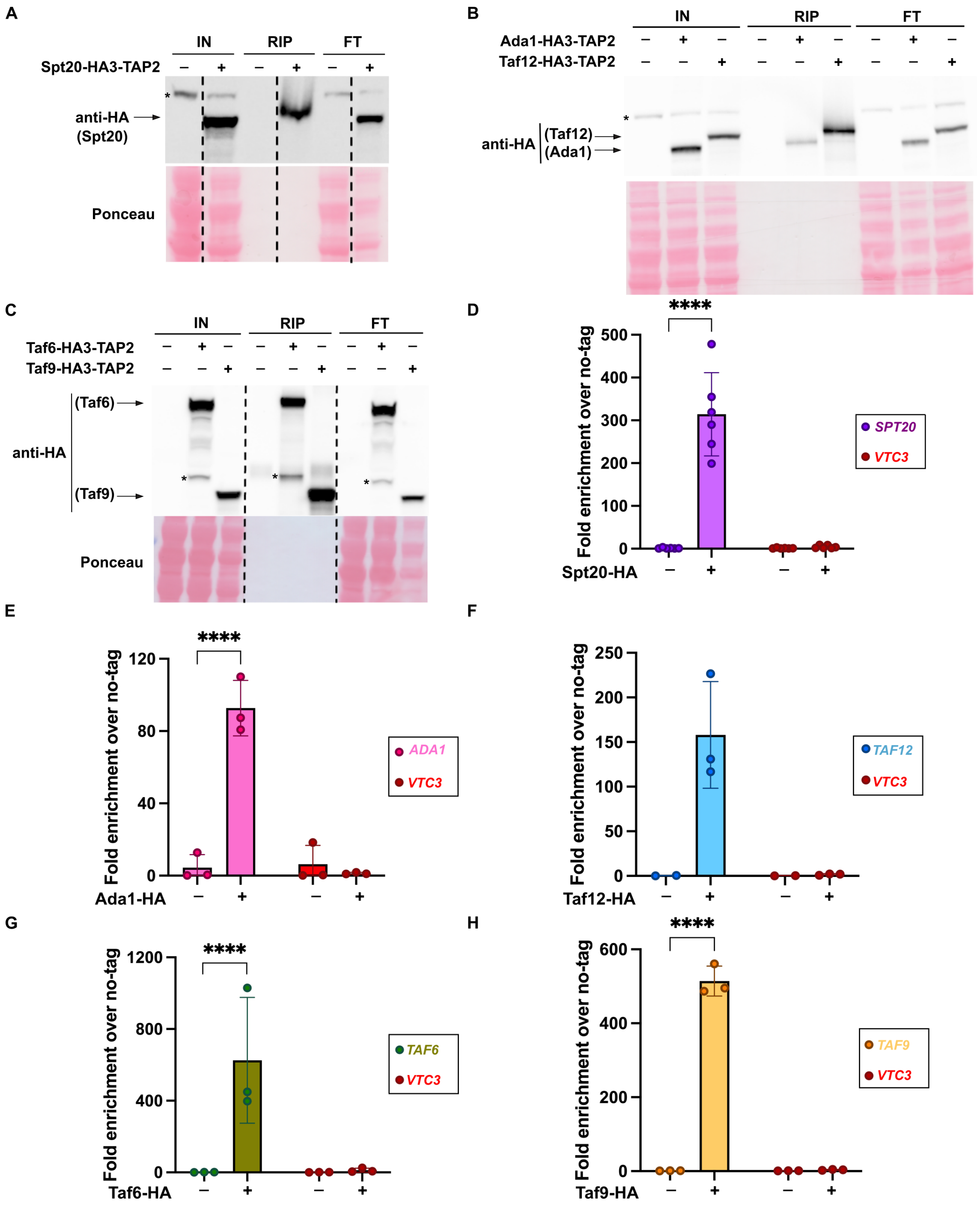


**Supplementary Figure S6. Immunoprecipitation and co-translational self-mRNA association controls for HA-tagged SAGA core subunits. (A–C)** Western blot controls for the anti-HA RIP-qPCR assays shown in Figure 7C–G, analyzing input (IN), immunoprecipitated (RIP), and flow-through (FT) fractions from untagged (−) and HA3-TAP2-tagged (+) strains: Spt20-HA3-TAP2 (A), Ada1-HA3-TAP2 and Taf12-HA3-TAP2 (B), and Taf6-HA3-TAP2 and Taf9-HA3-TAP2 (C). Membranes were probed with anti-HA to verify subunit expression and immunoprecipitation efficiency, and stained with Ponceau S to confirm protein loading and transfer. Asterisks (*) indicate non-specific cross-reacting bands, and vertical dashed lines in (A) and (C) indicate where intervening lanes from the same membrane were spliced out. **(D–H)** Anti-HA RIP-qPCR control analysis quantifying co-translational self-mRNA enrichment alongside the unrelated negative control *VTC3* in untagged (−) and HA3-TAP2-tagged (+) SAGA core subunit strains: Spt20-HA with *SPT20* mRNA (D), Ada1-HA with *ADA1* mRNA (E), Taf12-HA with *TAF12* mRNA (F), Taf6-HA with *TAF6* mRNA (G), and Taf9-HA with *TAF9*

mRNA (H). Individual data points represent independent biological replicates, with mean ± SD overlaid, expressed as fold enrichment ($2^{-\Delta\Delta Ct}$) relative to the mean $\Delta Ct$ of the untagged control. Individual replicates of this control are plotted to illustrate baseline variability. Statistical comparisons shown for the target SAGA core transcripts (− *vs.* +) in (D, E, G, H) were determined by two-way ANOVA on $\log_2$-transformed relative enrichment values followed by Tukey's multiple comparisons test (simple effects within each transcript), using a significance threshold of $\alpha = 0.01$ (****$p < 0.0001$); statistical analysis was omitted in (F) because $n = 2$ for the untagged (−) control. *VTC3* is plotted alongside in all panels as an unrelated negative control to illustrate baseline background.

**Supplementary Table S1. List of *S. cerevisiae* strains used in this work.**

| Name | Genotype | Source |
|---|---|---|
| BY4741 | *Mat a his3Δ1 leu2Δ0 met15Δ0 ura3Δ0* | Lab stock |
| BY4742 | *Mat α his3Δ1 leu2Δ0 lys2Δ0 ura3Δ0* | Lab stock |
| BY4745 | *Mat α his3Δ1 leu2Δ0 met15Δ0 ura3Δ0* | Lab stock |
| EVC36 | *BY4745 UBP8-NanoLuc::Hph* | Lazarewicz et al, 2024 |
| EVC40 | *BY4745 SGF11-NanoLuc::Hph* | Lazarewicz et al, 2024 |
| EVC43 | *BY4745 SUS1-NanoLuc::Hph* | Lazarewicz et al, 2024 |
| EVC44 | *BY4745 SGF73-NanoLuc::Hph* | Lazarewicz et al, 2024 |
| EVC43 | *BY4745 SGF11-LgBiT::Hph* | Le Boulch et al, 2020 |
| EVC50 | *BY4745 UBP8-LgBiT::Hph* | Le Boulch et al, 2020 |
| EVC53 | *BY4745 SGF11-LgBiT::Hph* | Le Boulch et al, 2020 |
| EVC57 | *BY4745 SUS1-LgBiT::Hph* | Le Boulch et al, 2020 |
| EVC76 | *BY4742 SGF73-GS-HAx3-CTAPx2::KanMX6+* | This work |
| EVC78 | *BY4742 SUS1-GS-HAx3-CTAPx2::KanMX6+* | This work |
| EVC80 | *BY4742 UBP8-GS-HAx3-CTAPx2::KanMX6+* | This work |
| EVC82 | *BY4742 SGF11-GS-HAx3-CTAPx2::KanMX6+* | This work |
| EVC90 | *SUS1-LgBiT::Hph UBP8-SmBiT::Nat* | This work |
| EVC95 | *BY4742 SGF73-GS-HAx3-CTAPx2::KanMX6+ sgf11Δ::NatMX6+* | This work |
| EVC97 | *BY4742 SUS1-GS-HAx3-CTAPx2::KanMX6+ sgf11Δ::NatMX6+* | This work |
| EVC98 | *BY4742 UBP8-GS-HAx3-CTAPx2::KanMX6+ sgf11Δ::NatMX6+* | This work |
| EVC100 | *BY4742 UBP8-GS-HAx3-CTAPx2::KanMX6+ sus1Δ::NatMX6+* | This work |
| EVC102 | *BY4742 SGF11-GS-HAx3-CTAPx2::KanMX6+ sus1Δ::NatMX6+* | This work |
| EVC104 | *BY4742 SGF73-GS-HAx3-CTAPx2::KanMX6+ ubp8Δ::NatMX6+* | This work |
| EVC106 | *BY4742 SUS1-GS-HAx3-CTAPx2::KanMX6+ ubp8Δ::NatMX6+* | This work |
| EVC108 | *BY4742 SGF11-GS-HAx3-CTAPx2::KanMX6+ ubp8Δ::NatMX6+* | This work |
| EVC109 | *BY4742 SGF11-GS-HAx3-CTAPx2::KanMX6+ sgf73Δ::NatMX6+* | This work |
| EVC112 | *BY4742 SGF73-GS-HAx3-CTAPx2::KanMX6+ sus1Δ::NatMX6+* | This work |
| EVC114 | *BY4742 SUS1-GS-HAx3-CTAPx2::KanMX6+ sgf73Δ::NatMX6+* | This work |
| EVC115 | *BY4742 SPT20-GS-HAx3-CTAPx2::KanMX6+* | This work |
| EVC117 | *BY4742 SUS1-GS-HAx3-CTAPx2::KanMX6+ SGF11-6xGly-FLAGx3::HphMX6+ ubp8Δ::NatMX6+* | This work |
| EVC119 | *BY4742 SUS1-GS-HAx3-CTAPx2::KanMX6+ SGF73-6xGly-FLAGx3::HphMX6+ ubp8Δ::NatMX6+* | This work |
| EVC121 | *BY4742 SUS1-GS-HAx3-CTAPx2::KanMX6+ SGF11-6xGly-FLAGx3::HphMX6+ sgf73Δ::NatMX6+* | This work |
| EVC123 | *BY4742 SUS1-GS-HAx3-CTAPx2::KanMX6+ SGF11-6xGly-FLAGx3::HphMX6+* | This work |
| EVC127 | *BY4742 SUS1-GS-HAx3-CTAPx2::KanMX6+ SGF73-6xGly-FLAGx3::HphMX6+* | This work |
| EVC131 | *BY4742 UBP8-GS-HAx3-CTAPx2::KanMX6+ SGF73-6xGly-FLAGx3::HphMX6+* | This work |
| EVC133 | *BY4742 UBP8-GS-HAx3-CTAPx2::KanMX6+ SGF73-6xGly-FLAGx3::HphMX6+ sgf11Δ::NatMX6+* | This work |
| EVC135 | *BY4742 SGF11-GS-HAx3-CTAPx2::KanMX6+ SGF73-6xGly-FLAGx3::HphMX6+* | This work |
| EVC137 | *BY4742 SGF11-GS-HAx3-CTAPx2::KanMX6+ SGF73-6xGly-FLAGx3::HphMX6+ sus1Δ::NatMX6+* | This work |
| EVC138 | *BY4742 SGF11-GS-HAx3-CTAPx2::KanMX6+ SGF73-6xGly-FLAGx3::HphMX6+ ubp8Δ::NatMX6+* | This work |
| EVC139 | *BY4742 UBP8-GS-HAx3-CTAPx2::KanMX6+ SGF73-6xGly-FLAGx3::HphMX6+ sus1Δ::NatMX6+* | This work |
| EVC141 | *BY4742 ADA1-GS-HAx3-CTAPx2::KanMX6+* | This work |
| EVC145 | *BY4742 SUS1-GS-HAx3-CTAPx2::KanMX6+ SGF73-6xGly-FLAGx3::HphMX6+ sgf11Δ::NatMX6+* | This work |

**Supplementary Table S1 (cont.). List of *S. cerevisiae* strains used in this work.**

| Name | Genotype | Source |
|---|---|---|
| EVC146 | *BY4745 UBP8-NanoLuc::Hph sgf11Δ::KanMX6+* | This work |
| EVC148 | *BY4745 SUS1-NanoLuc::Hph sgf11Δ::KanMX6+* | This work |
| EVC150 | *SUS1-LgBiT::Hph UBP8-SmBiT::Nat sgf11Δ::KanMX6+* | This work |
| EVC153 | *BY4742 sgf73Δ::NatMX6+* | This work |
| EVC155 | *BY4742 sus1Δ::NatMX6+* | This work |
| EVC160 | *BY4745 SGF11-NanoLuc::Hph sus1Δ::KanMX6+* | This work |
| EVC162 | *BY4742 sgf11Δ::NatMX6+* | This work |
| EVC164 | *BY4742 ubp8Δ::NatMX6+* | This work |
| EVC168 | *BY4745 SUS1-NanoLuc::Hph ubp8Δ::KanMX6+* | This work |
| EVC170 | *BY4745 SGF11-NanoLuc::Hph ubp8Δ::KanMX6+* | This work |
| EVC172 | *BY4745 UBP8-NanoLuc::Hph sus1Δ::KanMX6+* | This work |
| EVC174 | *BY4742 TAF12-GS-HAx3-CTAPx2::KanMX6+* | This work |
| EVC178 | *BY4745 SGF73-NanoLuc::Hph sus1Δ::KanMX6+* | This work |
| EVC180 | *BY4745 SGF73-NanoLuc::Hph ubp8Δ::KanMX6+* | This work |
| EVC182 | *BY4745 SGF73-NanoLuc::Hph sgf11Δ::KanMX6+* | This work |
| EVC184 | *BY4745 UBP8-NanoLuc::Hph sgf73Δ::KanMX6+* | This work |
| EVC186 | *BY4745 SUS1-NanoLuc::Hph sgf73Δ::KanMX6+* | This work |
| EVC188 | *BY4742 SUS1-GS-HAx3-CTAPx2::KanMX6+ UBP8-MYCx13::NatMX6+* | This work |
| EVC190 | *SGF11-LgBiT::Hph SGF73-SmBiT::Nat* | This work |
| EVC192 | *SUS1-LgBiT::Hph SGF73-SmBiT::Nat* | This work |
| EVC198 | *SUS1-LgBiT::Hph UBP8-SmBiT::Nat sgf73Δ::KanMX6+* | This work |
| EVC199 | *SGF11-LgBiT::Hph SGF73-SmBiT::Nat sus1Δ::KanMX6+* | This work |
| EVC201 | *SGF11-LgBiT::Hph SGF73-SmBiT::Nat ubp8Δ::KanMX6+* | This work |
| EVC202 | *SUS1-LgBiT::Hph SGF73-SmBiT::Nat sgf11Δ::KanMX6+* | This work |
| EVC204 | *SUS1-LgBiT::Hph SGF73-SmBiT::Nat ubp8Δ::KanMX6+* | This work |
| EVC205 | *UBP8-LgBiT::Hph SGF11-SmBiT::Nat* | This work |
| EVC207 | *UBP8-LgBiT::Hph SGF11-SmBiT::Nat sus1Δ::KanMX6+* | This work |
| EVC209 | *UBP8-LgBiT::Hph SGF11-SmBiT::Nat sgf73Δ::KanMX6+* | This work |
| EVC212 | *UBP8-LgBiT::Hph SGF73-SmBiT::Nat* | This work |
| EVC213 | *SGF11-LgBiT::Hph SUS1-SmBiT::Nat* | This work |
| EVC215 | *UBP8-LgBiT::Hph SGF73-SmBiT::Nat sus1Δ::KanMX6+* | This work |
| EVC216 | *UBP8-LgBiT::Hph SGF73-SmBiT::Nat sgf11Δ::KanMX6+* | This work |
| EVC218 | *SGF11-LgBiT::Hph SUS1-SmBiT::Nat ubp8Δ::KanMX6+* | This work |
| EVC220 | *SGF11-LgBiT::Hph SUS1-SmBiT::Nat sgf73Δ::KanMX6+* | This work |
| EVC222 | *BY4742 SUS1-GS-HAx3-CTAPx2::KanMX6+ UBP8-MYCx13::NatMX6+ sgf73Δ::HphMX6+* | This work |
| EVC223 | *BY4742 SUS1-GS-HAx3-CTAPx2::KanMX6+ UBP8-MYCx13::NatMX6+ sgf11Δ::HphMX6+* | This work |
| EVC225 | *BY4742 SGF11-GS-HAx3-CTAPx2::KanMX6+ UBP8-MYCx13::NatMX6+* | This work |
| EVC234 | *BY4742 SGF11-GS-HAx3-CTAPx2::KanMX6+ UBP8-MYCx13::NatMX6+ sus1Δ::HphMX6+* | This work |
| EVC236 | *BY4742 SGF11-GS-HAx3-CTAPx2::KanMX6+ UBP8-MYCx13::NatMX6+ sgf73Δ::HphMX6+* | This work |
| EVC244 | *BY4742 TAF6-GS-HAx3-CTAPx2::KanMX6+* | This work |
| EVC246 | *BY4742 TAF9-GS-HAx3-CTAPx2::KanMX6+* | This work |

**Supplementary Table S1 (cont.). List of *S. cerevisiae* strains used in this work.**

| Name | Genotype | Source |
|---|---|---|
| EVC248 | *BY4742 UBP8-MYCx13::NatMX6+* | This work |
| EVC274 | *BY4742 UBP8-GS-HAx3-CTAPx2::KanMX6+ sgf73Δ::NatMX6+* | This work |

**Supplementary Table S2. List of oligonucleotides used in this work.**

| Gene / plasmid | Name | Description | Sequence |
|---|---|---|---|
| *ADA1* | EVO47 | C-terminal tagging 5'-end / pFA6a | GTACAAGGCATGAATTAAATTGGTTAATTAAAGGAATCTTAACGGAAGATCGGATCCCCGGGTTAATTAA |
| *ADA1* | EVO48 | C-terminal tagging 3'-end / pFA6a | TAATCTTTCAATATAATTACAACATACCGCATACACACACTTTTTATACAGAATTCGAGCTCGTTTAAAC |
| *ADA1* | EVO49 | Check C-terminal tagging 5'-end | AGCAGCATTACAAAGGCTGG |
| *ADA1* | EVO50 | Check C-terminal tagging 3'-end | TAAGCTCGGTCACATTATCG |
| *ADA1* | EVO377 | RIP-qPCR | AAACCCTCTTGGGAGAAACG |
| *ADA1* | EVO378 | RIP-qPCR | AATGAAGCCTCTTTTGCCGG |
| *pFA6a* | EVO21 | Check C-terminal tagging 5'-end or KO 5'-end | TTAATTAACCCGGGGATCCG |
| *pFA6a* | EVO22 | Check C-terminal tagging 3'-end or KO 3'-end | GTTTAAACGAGCTCGAATTC |
| *SGF11* | EVO204 | C-terminal tagging 5'-end / pFA6a | ACAGACTAGCAGCTCATTTACAGAGATGTTTGAGTAGGGGTGCTAGACGTCGGATCCCCGGGTTAATTAA |
| *SGF11* | EVO205 | C-terminal tagging 3'-end or KO 3'-end / pFA6a | GCTAAAATCTGTCTGTGCCTTTTCAATTACCCATAAACCACCACCTAGTGGAATTCGAGCTCGTTTAAAC |
| *SGF11* | EVO206 | Check C-terminal tagging 5'-end | TACAGAAGCAACAAGAGTCC |
| *SGF11* | EVO207 | Check C-terminal tagging 3'-end or KO 3'-end | TTGGCATCATATATGGGAGG |
| *SGF11* | EVO309 | RIP-qPCR / RT-qPCR | TCCAGGACATTGTAGCTCGG |
| *SGF11* | EVO310 | RIP-qPCR / RT-qPCR | ACTGAGAGGACTCTTGTTGC |
| *SGF11* | EVO358 | KO 5'-end / pFA6a | GTAACTTAGCGTTTCTTCCGTTGAGCAGAGCAGTATAATAATCCTTCAAATTCTTTAGGTTACGGGGTTTTCCTGTTGCGCGGATCCCCGGGTTAATTAA |
| *SGF11* | EVO361 | Check KO 5'-end | GCAGTGATTTGCTTATTTTCT |
| *SGF73* | EVO348 | C-terminal tagging 5'-end / pFA6a | TTGAAGTAGGTATTGGAAATTCTGTGAACCCCTACAATGGCAGAATAAATCGGATCCCCGGGTTAATTAA |
| *SGF73* | EVO349 | C-terminal tagging 3'-end or KO 3'-end / pFA6a | ATTTTTTATTACTCACTTCGTGAACATGCTGGATAACGTGCATGATTCAAGAATTCGAGCTCGTTTAAAC |
| *SGF73* | EVO89 | Check C-terminal tagging 5'-end | AACCTCATGACGATGAAAGC |
| *SGF73* | EVO90 | Check C-terminal tagging 3'-end or KO 3'-end | AAATAGCGTGACTTGTCCAC |
| *SGF73* | EVO305 | RIP-qPCR / RT-qPCR | GCACAACTCATTGAAGATCC |
| *SGF73* | EVP306 | RIP-qPCR / RT-qPCR | CTCATCTCTTGGATCTGTGC |
| *SGF73* | EVO350 | KO 5'-end / pFA6a | TTAATTTACCAAAACGGAATAAACCAAAAAAATAAGAATAGTGAACACACAAGAGAAGCGCAAAAGAGTAAAGAGCTAAACGGATCCCCGGGTTAATTAA |
| *SGF73* | EVO351 | Check KO 5'-end | GAACCTTTTTGATATCCAGC |
| *SPT20* | EVO91 | C-terminal tagging 5'-end / pFA6a | CTAAGAAGAAACAGAGCGCAAGTAGCACGCCCTCTTCTACTACAATGTCACGGATCCCCGGGTTAATTAA |
| *SPT20* | EVO360 | C-terminal tagging 3'-end / pFA6a | ATATATATATATATATATATATAAGGAATGATAACTCTATTTAAGTAGAGAATTCGAGCTCGTTTAAAC |
| *SPT20* | EVO93 | Check C-terminal tagging 5'-end | AAACGGAGTGGTAACAATGC |
| *SPT20* | EVO94 | Check C-terminal tagging 3'-end | TCTTTTGCTTGAGCACAGAGG |
| *SPT20* | EVO361 | RIP-qPCR | ATGCCAATGAAGGTACTAGG |
| *SPT20* | EVO362 | RIP-qPCR | AAATTGAGTCTGCATCTTCC |
| *SUS1* | EVO352 | C-terminal tagging 5'-end / pFA6a | CCGTTTTAAAGCAAATAAGGGAATTTCTTGAAGAGATTGTAGATACACAACGGATCCCCGGGTTAATTAA |
| *SUS1* | EVO353 | C-terminal tagging 3'-end or KO 3'-end / pFA6a | CGATGAGCATATGTAATAATATTGGGAATTAAGGTGCATTTTCGTATCCTGAATTCGAGCTCGTTTAAAC |
| *SUS1* | EVO202 | Check C-terminal tagging 5'-end | ATCCACCGTAGAACCCAAAGC |
| *SUS1* | EVO203 | Check C-terminal tagging 3'-end or KO 3'-end | AGGTGGGTAACGTGAATTAGG |
| *SUS1* | EVO307 | RIP-qPCR / RT-qPCR | ATTTCAAACGAACTAAAAGC |
| *SUS1* | EVO308 | RIP-qPCR / RT-qPCR | CTAATGCTTTGGGTTCTACG |
| *SUS1* | EVO354 | KO 5'-end / pFA6a | AAAAAATGAAAATTGCCATAGGCATGGCTTGACCAACACATGCGACAAAATCAGAAGTAACAATTCTGGCCTTCACTCCACGGATCCCCGGGTTAATTAA |
| *SUS1* | EVO355 | Check KO 5'-end | ACCTCTTCCTCCATGAAGACC |
| *TAF12* | EVO392 | C-terminal tagging 5'-end / pFA6a | CAGCTGCAAAAAACAATGGAAACAATGTTGCAAGCTTGAATACAAAAAAACGGATCCCCGGGTTAATTAA |
| *TAF12* | EVO393 | C-terminal tagging 3'-end / pFA6a | TAACTTTAGGAATTTTTTCTGTGGAAGAGTGGAGAATGTGGGTAACATAAGAATTCGAGCTCGTTTAAAC |

**Supplementary Table S2 (cont.). List of oligonucleotides used in this work.**

| Gene / plasmid | Name | Description | Sequence |
|---|---|---|---|
| *TAF12* | EVO394 | Check C-terminal tagging 5'-end | TGGAATATTAGGATTCCTGG |
| *TAF12* | EVO395 | Check C-terminal tagging 3'-end | GAGAGCGTAGTTATGATGGG |
| *TAF12* | EVO375 | RIP-qPCR / RT-qPCR | AGCCGCAAAGATCAAACAGG |
| *TAF12* | EVO376 | RIP-qPCR / RT-qPCR | ATTCTTGGTTCTCTTCCGGG |
| *UBP8* | EVO208 | C-terminal tagging 5'-end / pFA6a | TTTTAAAGGAACAGGCATATTTATTATTCTACACCATTCGTCAAGTAAAT CGGATCCCCGGGTTAATTAA |
| *UBP8* | EVO209 | C-terminal tagging 3'-end or KO 3'-end / pFA6a | CTTTTTCTTCTTTTTTGTTTTATTATTATTGTTGAATGCTATTTGCTGAA GAATTCGAGCTCGTTTAAAC |
| *UBP8* | EVO210 | Check C-terminal tagging 5'-end | TCTGAAAATGGCAAGGTTCC |
| *UBP8* | EVO211 | Check C-terminal tagging 3'-end or KO 3'-end | ACTCACTTAGTTTTCCAGCC |
| *UBP8* | EVO311 | RIP-qPCR / RT-qPCR | TGGATGTTGGAACCATTCGC |
| *UBP8* | EVO312 | RIP-qPCR / RT-qPCR | TCTTTGTGCACACGTCGTCC |
| *UBP8* | EVO356 | KO 5'-end / pFA6a | CGGATCAAAAGGGAAAACTACAGCTTCGGTCCTCGTCGTCCTACTTGAAACCCTG CTTTTTTTATTTGTTATTAATAATTCGGATCCCCGGGTTAATTAA |
| *UBP8* | EVO357 | Check KO 5'-end | AGTCCATTTCTTTCAACTGG |
| *TAF6* | EVO420 | C-terminal tagging 5'-end / pFA6a | TGAAAAGAGACGACGCAAAAGAATTAATTAGTGCGATATTTTTTGGCGAACGGA TCCCCGGGTTAATTAA |
| *TAF6* | EVO421 | C-terminal tagging 3'-end / pFA6a | AAACTGTTATATAACATTACCGTTGTACTATTAGTATAAACCGATTCTCAGAATTC GAGCTCGTTTAAAC |
| *TAF6* | EVO422 | Check C-terminal tagging 5'-end | CGATGACTCCAATCCAACTA |
| *TAF6* | EVO423 | Check C-terminal tagging 3'-end | GCTCCAGAATCCACATAGAA |
| *TAF6* | EVO428 | RIP-qPCR | ACAGACGATGTATCGAAGGC |
| *TAF6* | EVO429 | RIP-qPCR | AAAGTTGGTAGACGAGGCAC |
| *TAF9* | EVO424 | C-terminal tagging 5'-end / pFA6a | AATATTGTCTCACAGCGAAGGAGTGGGACCTTGAGGACCCAAAGTCTATGCGGATCCCCGG GTTAATTAA |
| *TAF9* | EVO425 | C-terminal tagging 3'-end / pFA6a | TCTTCCGGAAGAACTTGATTTAGTGTACTTATGATATATAAGCGGTGCATGAATT CGAGCTCGTTTAAAC |
| *TAF9* | EVO426 | Check C-terminal tagging 5'-end / RIP-qPCR | GGACCAAGTACCGCTACAAT |
| *TAF9* | EVO427 | Check C-terminal tagging 3'-end | GCCTATCACGTCTTCTTTCA |
| *TAF9* | EVO430 | RIP-qPCR | AACATGAGCTCCTTAGGTGC |

**Suplementary references**


1. Le Boulch M, Brossard A, Le Dez G, Léon S, Rabut G. Sensitive detection of protein ubiquitylation using a protein fragment complementation assay. J Cell Sci. 2020 Jun 15;133(12).
2. Lazarewicz N, Le Dez G, Cerjani R, Runeshaw L, Meurer M, Knop M, et al. Accurate and sensitive interactome profiling using a quantitative protein-fragment complementation assay. Cell Reports Methods. 2024 Oct;4(10):100880.
